# Transient activation of the UPR^ER^ is an essential step in the acquisition of pluripotency during reprogramming

**DOI:** 10.1101/472787

**Authors:** Milos S. Simic, Erica Moehle, Robert T. Schinzel, Franziska Lorbeer, Damien Jullié, Jonathan J. Halloran, Kartoosh Heydari, Melissa Sanchez, Dirk Hockemeyer, Andrew Dillin

## Abstract

Somatic cells can be reprogrammed into pluripotent stem cells by the forced expression of the OCT4, SOX2, KLF4 and c-MYC transcription factors. This process requires the reshaping of not only epigenetic landscapes, but the global remodeling of cell identity, structure, and function including such basic processes of metabolism and organelle form and function. Cellular reprogramming is a stochastic process with only a marginally measureable fraction of cells successfully crossing these, and many other, cellular epitomes to acquire the fully pluripotent state. We hypothesize that this variation is due, in part, by variable regulation of the proteostasis network and its influence upon the protein folding environment within cells and their organelles upon the remodeling process. We find that the endoplasmic reticulum unfolded protein response (UPR^ER^), the heat-shock response (HSR) and the mitochondrial unfolded protein response (UPR^mt^), which monitor and ensure the quality of the proteome of, respectively, the ER, the cytosol and the mitochondria during stress, are activated during cellular reprogramming. Particularly, we find that the UPR^ER^ is essential for reprograming, and ectopic, transient activation of the UPR^ER^, either genetically or pharmacologically, enhances the success of cells to reach a pluripotent state. Finally, and most revealing, we find that stochastic activation of the UPR^ER^ can predict the reprogramming efficiency of naïve cells. The results of these experiments indicate that the low efficiency and stochasticity of cellular reprogramming is partly the result of the inability to initiate a proper ER stress response for remodeling of the ER and its proteome during the reprogramming process. The results reported here display only one aspect of the proteostasis network and suggest that proper regulation of many more components of this network might be essential to acquire the pluripotent state.

## Introduction

Reprogramming of somatic cells into induced pluripotent stem cells (iPSCs) highlights the remarkable plasticity found within cells and provides an incredible potential for cell biology and regenerative medicine (1). Cellular reprogramming can be achieved by the forced expression of OCT4, SOX2, KLF4 and c-MYC, transcription factors with a wide range of target genes (2). However, the success of cellular reprogramming of human cells is extremely low, ranging from.0001% to.1%. The mechanisms that drive the variability and stochastic nature of reprogramming are enigmatic and pose one of the major hurdles in the reprogramming process (3, 4). Therefore, a better understanding of the mechanisms underlying reprogramming is necessary to improve this process (5).

It is clear that genome integrity and epigenetic rewiring are central tenants of the reprogramming process and could explain much of the variation within the acquisition of the pluripotent state. However, as the somatic cell transitions into a new identity with changes in epigenetics wiring, the constituents and quality of its sub-cellular organelles are also undergoing massive re-wiring and are under selective pressure to ensure a pristine proteome of the resulting, immortal iPSC. Inheritance of faulty proteins and organelles provide challenges upon a cell driving towards immortality and pluripotency. Therefore, the stress during this process is not only confined within the nucleus, but emanates throughout the cell and subcellular organelles. To ensure a proper balance of proteome function and organelle integrity, a delicate network exists that monitors and responds to challenges within the proteomes of sub-cellular organelles, known as the proteostasis network. Within the proteostasis network, key stress responses, such as the UPR^ER^, which monitors the integrity of the endoplasmic reticulum, the UPR^mt^, which monitors mitochondrial quality and the HSR, which predominantly interrogates the cytoplasm, govern and dictate proteome fidelity and organelle function (6).

Secreted and membrane-bound proteins are synthesized in the endoplasmic reticulum (ER) and represent up to one third of the total proteome produced by cells. Increased protein synthesis, cell differentiation, tissue development, senescence, DNA damage and many other stressors, disrupt ER homeostasis and activate the UPR^ER^ (7). Three ER-resident transmembrane proteins sense the protein folding state in the ER lumen and transduce this information using parallel and distinctive signal transduction mechanisms: ATF6 (activating transcription factor 6), PERK (double-stranded RNA-activated protein kinase (PRK)-like ER kinase) and IRE1 (inositol requiring enzyme 1) (7). During stress, IRE1 converges on the X-box binding protein 1 transcription factor, XBP1, causing its cytoplasmic splicing to create the *XBP1s* mRNA that can be translated and incorporated into the nucleus to regulate hundreds of genes required for ER protein folding and morphology (8, 9). PERK functions to decrease global translation by phosphorylation of eIF2α (*10*) while specifically increasing translation of the transcription factor ATF4 (*11*). Furthermore, ATF6 is shuffled from the ER to the Golgi where two Golgi-resident proteases cleave it, releasing its cytosolic DNA-binding domain that enters the nucleus and activates target genes (12).

Cellular reprogramming causes a dramatic change in cell morphology and imposes the remodeling of many organelles such as mitochondria (*13).* We therefore hypothesized that cellular reprogramming should restructure the ER and require the UPR^ER^. Furthermore, the UPR^ER^ presents stochastic variation amongst isogenic cell populations with some cells mounting a robust response and others feebly attempting induction. We further speculated that the UPR^ER^ might not only play a pivotal role during reprogramming, but could also explain its stochastic nature and could predict, at least in part, this inherent stochasticity.

## Results

### Cellular reprogramming activates the UPR^ER^, HSR and UPR^mt^

During stress, the transcription of central regulators of the proteostasis network are increased as well as their downstream targets (6). We analyzed the canonical downstream transcriptional targets of the UPR^ER^ (HSPA5 and GRP94), HSR (HSPA1A) and UPR^mt^ (GRP75) during reprogramming of neonatal fibroblasts and found that transcriptional targets of each response were increased compared to cells not undergoing reprogramming (Fig. 1A). This observation was extended to reprogramming of neonatal keratinocytes as well (Fig. S1A). During the reprogramming process, the 4 reprogramming factors are delivered by viral infection. To exclude the possibility that the UPR^ER^ is induced by the use of a viral delivery system, we also used an episomal delivery system of the reprogramming factors and found similar activations of the HSR, UPR^ER^ and UPR^mt^ (Fig. S1B). To corroborate the mRNA levels, we analyzed HSPA5, GRP94, HSPA1A and GRP75 protein levels and found that they too were increased (Fig. 1B and S1C). The differences in protein levels between GFP Day 3 (D3) and GFP Day 6 (D6) is due to the time at which cells were moved to iPS reprogramming media on day 4. Therefore each comparison was normalized to its respective control GFP. The activation of the UPR^ER^ and HSR were to that found in cells undergoing an ER stress, tunicamycin, or a heat-shock, 42°C (Fig. S1D and S1E). We also confirmed by both mRNA and protein levels that overexpression of GFP did not activate the stress pathways (Fig. 1B, S1F and S1G).

**Fig. 1:**
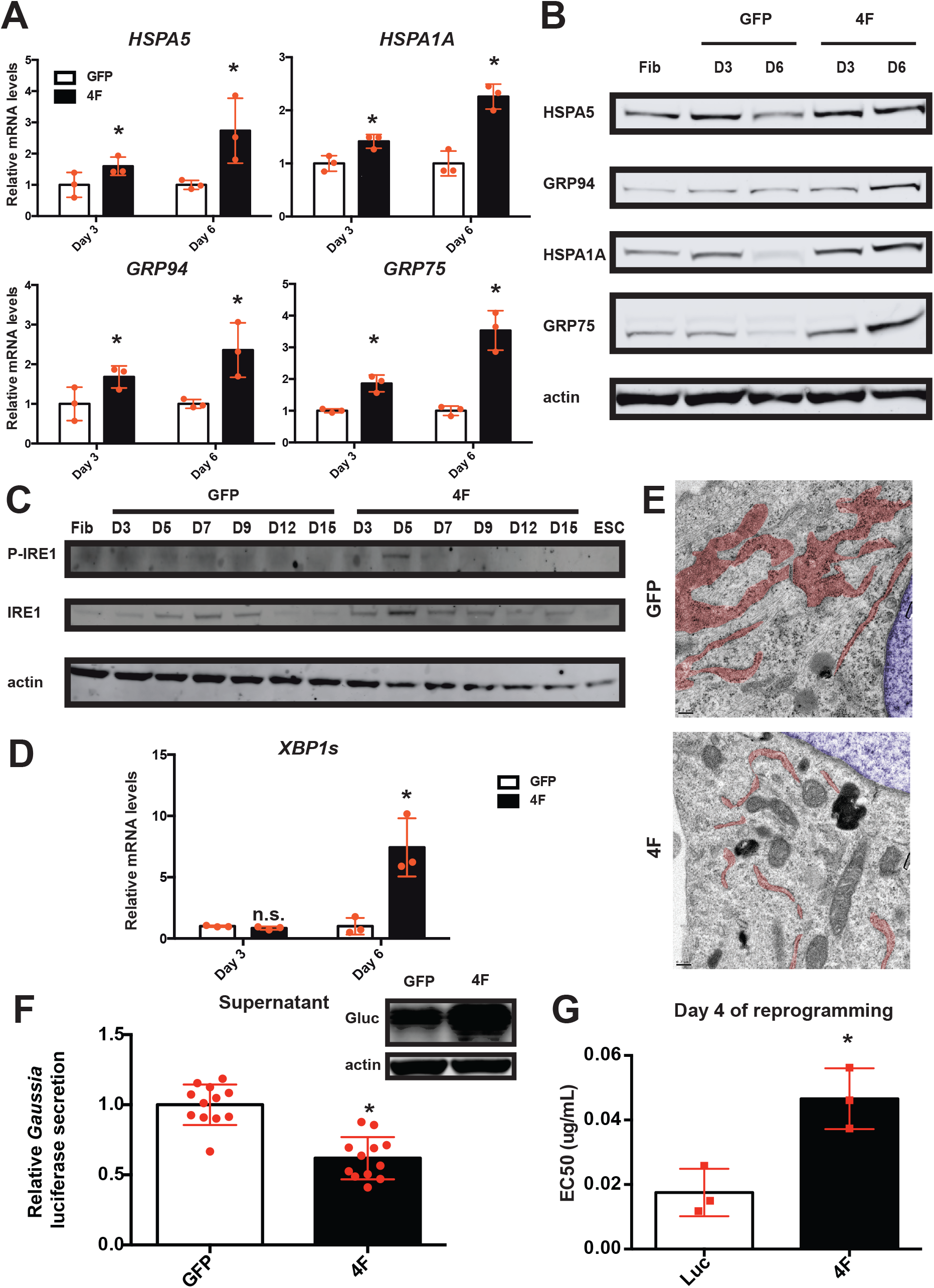
The three major unfolded protein responses are activated during cellular reprogramming. (A) Relative mRNA levels of the main effectors of the UPR^ER^ (HSPA5 and GRP94), HSR (HSPA1A) and UPR^mt^ (GRP75) relative to *GAPDH* determined by qRT-PCR (n=3, average +/− SD). GFP control was set to 1 for each day. (B) Western blot analysis of the main effectors of the UPR^ER^ (HSPA5 and GRP94), HSR (HSPA1A) and UPR^mt^ (GRP75). (C) Time course reprogramming western blot analysis of P-IRE1 and IRE1. (D) Relative mRNA levels of the spliced form of *XBP1* relative to *GAPDH* determined by qRT-PCR (n=3, average +/− SD). GFP control was set to 1 for each day. (E) Electron microscopy of day 4 reprogramming fibroblasts and GFP control, scale bar = 0.2 μm. Pseudo-colors blue and red mark respectively the nucleus and the ER. (F) Secretion capability of the ER measured by luciferase activity secreted in the media (n=12, average +/− SD) and western blot analysis of the *Gaussia* luciferase. (G) Sensitivity to tunicamycin treatment determined by EC50 measurement at day 4 of reprogramming of fibroblast-like cells (n=3, average +/− SD). * indicates statistical difference (p-value<0.05) using an unpaired two-tailed t-test, n.s. indicates statistical nonsignificance.

Because of the important role of the UPR^ER^ in stem cells and during differentiation (*14),* we decided to further characterize its activation during cellular reprogramming. We analyzed the phosphorylated state of IRE1 and PERK, modifications indicative of ER stress, and found that both were highly phosphorylated during the reprogramming process (Fig. 1C and S2A). Interestingly, in all cases, we observed a transient upregulation of the UPR^ER^ that was not prolonged or extended after the acquisition of pluripotency. Phosphorylation of IRE1 leads to the cytosolic splicing of XBP1 mRNA. Consistent with activation of IRE1, we observed increased spliced mRNA of XBP1 in both fibroblasts (Fig. 1D) and keratinoyctes undergoing reprogramming (Fig. S2B). The mRNA levels of CHOP, a canonical downstream target of the PERK pathway, was also increased in both fibroblasts and keratinoyctes (Fig. S2C). Finally we tested the activation of the third branch of the UPR^ER^ pathway, the transcriptional activation of ATF6 (15). We found ATF6 mRNA levels in both fibroblasts and keratinocytes were increased during cellular reprogramming (Fig. S2D).

The ER is composed of an orchestrated architecture that can be dynamic to include tubular geometry fused with undulant sheets. By electron microscopic (EM) analysis, the ER, pseudo colored in red, of cells undergoing reprogramming appears largely tubular, lacking sheet structures (Fig. 1E). The network and the high branching aspect seen in control cells are lost during reprogramming. It appears that the volume of the ER is decreased as well. In fact, the ER of cells undergoing reprogramming resembles cells treated with the ER stressor, tunicamycin (Fig. S2E). Molecularly, levels of Reticulon 4 (a marker of tubular ER) were increased and CLIMP-63 (a marker of cisternae/sheets) was decreased (16) during reprogramming, consistent with the EM analysis revealing tubular ER structures and few sheet structures (Fig. S2A).

Tubular ER morphology is associated with impaired secretory capacity. We tested the secretion capacity of cells undergoing reprogramming by following the secretion of the exogenously expressed humanized *Gaussia* luciferase protein (Gluc) (17). We collected the supernatant of cells undergoing reprogramming and observed a dramatic reduction in secreted Gluc. Importantly, the reduced secretion of Gluc during reprogramming was not due to decreased expression of the Gluc transgene during the reprogramming process (Fig. 1F).

Consistent with increased ER stress, morphological remodeling of the ER and reduced ER secretory function during the reprogramming process, we also found that cells undergoing reprogramming were more resistant to exogenous ER stress than control cells. Using a dose-survival curve for cells grown in the presence of tunicamycin, we found that cells undergoing reprogramming were more protected than cells not attempting to acquire pluripotency (Fig. 1G). In sum, ER stress, morphology and function are dramatically altered during the cellular reprogramming process and it appears to be transient and not retained in the ensuing pluripotent cell.

### Advanced states of reprogramming positively correlate with UPR^ER^ activation

Intrigued by the findings that the ER undergoes profound changes as a cell transitions from a basic, unilateral fate to one that is expansive and pluripotent, we began to query the major driver of ER remodeling and stress to understand what role, if any, did the UPR^ER^ play in cellular reprogramming. To decipher the role of the UPR^ER^ during reprogramming and test if it could be a limiting factor (i.e. essential) for successful reprogramming, we created somatic cells that contained a visible marker of UPR^ER^ induction. Briefly, we followed induction of the endogenous UPR^ER^ target gene HSPA5 by fusing eGFP onto its C-terminus. Using transcription activator-like effector nuclease (TALENs) mediated genome editing, we inserted eGFP to the last amino acid of HSPA5 in H9 embryonic stem cells (ESCs) (Fig. S3A). Successful targeting was confirmed by southern blot (Fig. 2A) and western blot analysis (Fig. 2B) as the predicted HSPA-GFP fusion protein is recognized by both GFP and HSPA5 antibodies. No other GFP specific bands were observed suggesting that any potential off-target integrations were not translated. The proper integration was further confirmed by sequencing of the targeted locus (Fig. S3A). The HSPA5-GFP cell line was then differentiated into somatic fibroblast-like cells (*18*). The resulting somatic cells were then used for cellular reprogramming to assess how the UPR^ER^ responded during cellular reprograming. As a control for reprogramming experiments, the HSPA5-GFP somatic cells responded faithfully to ER stress caused by tunicamycin, showing robust GFP fluorescence detectable by fluorescence microscopy (data not shown), protein levels (Fig. 2B) and fluorescent activated cell analysis (Fig. 2C). Importantly, the induction was reversible. After removal of tunicamycin, GFP levels decreased over time in these reporter cells (Fig. 2C), indicating that the reporter faithfully portrayed ER stress induction and not overt cellular damage.

**Fig. 2:**
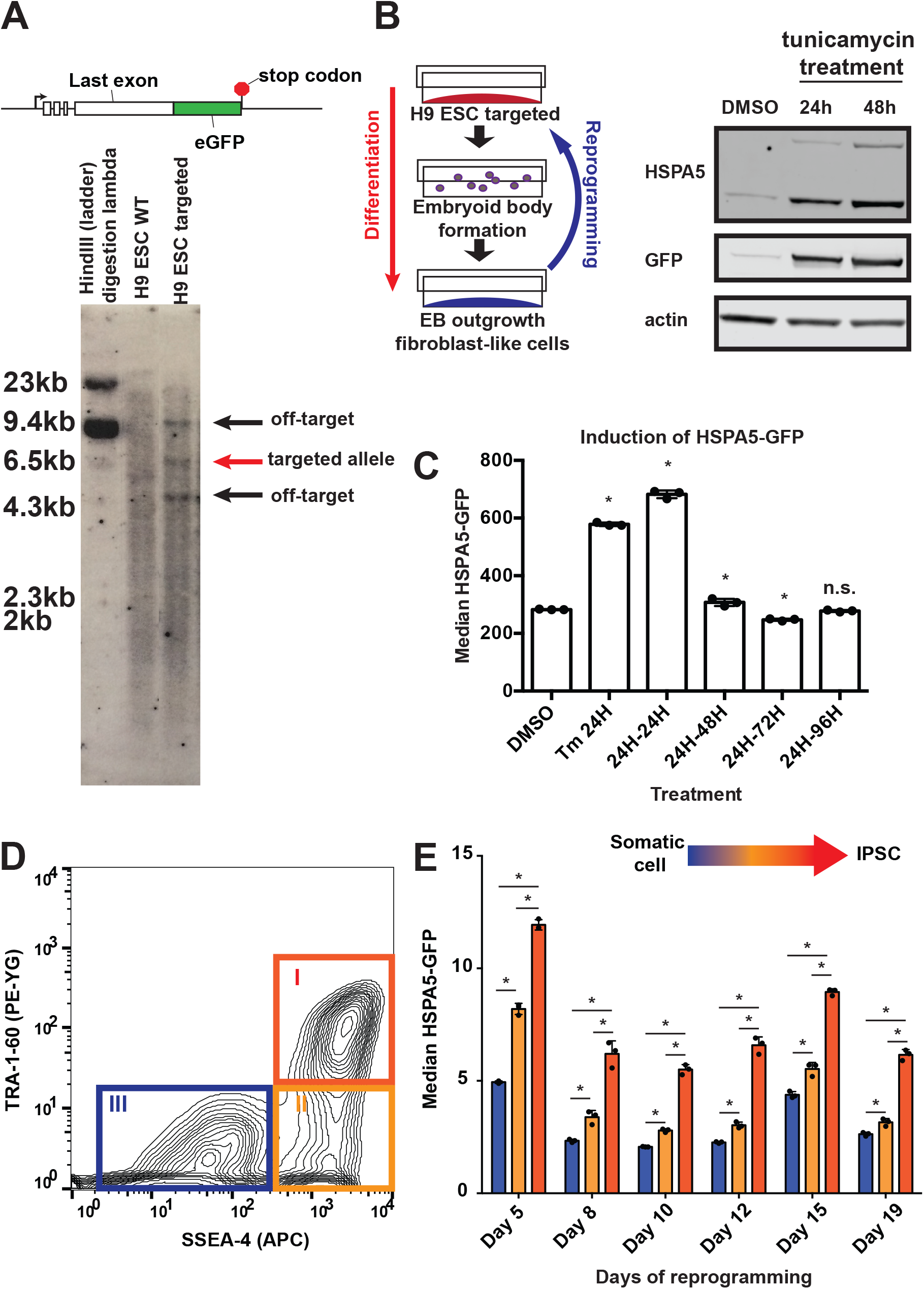
Advanced state of reprogramming positively correlates with higher UPR^ER^ activation. (A) Schematic of the genome editing strategy and southern blot using a GFP probe. The red arrow shows the expected size of the targeted allele while the black arrows show two off-target integrations. (B) Schematic of the fibroblast-like cells differentiation protocol (left panel) and western blot of HSPA5, GFP and actin showing the dynamical induction of the reporter line after the addition of 0.1μg/mL tunicamycin. The predicted HSPA5-GFP fusion band was targeted by both GFP and HSPA5 antibodies using dual channel imaging with the Odyssey^®^ CLx Imaging System confirming the correct targeting. Only a single intense specific GFP band was observed suggesting the off-targets integrations are not translated (right panel). (C) Median HSPA5-GFP levels analyzed by flow cytometry upon 0.1μg/mL tunicamycin treatment during 24h and after removal (n=3, average +/− SD). * indicates statistical difference (p-value<0.05) using Dunnett’s multiple comparison test to the DMSO control. (D) Flow cytometry analysis of fibroblast-like HSPA5-GFP cells at day 8 of reprogramming stained with SSEA-4 and TRA-1-60 surface markers. I, II, III represent the different cell states of reprogramming. (E) Median HSPA5-GFP of the different cell states (I, II, III) during reprogramming (n=3, average +/− SD). * indicates statistical difference (p-value<0.05) using Newman-Keuls multiple comparison test between all the conditions.

Equipped with a reliable, live cell marker for ER stress, we now needed to couple it to molecular signatures of the process of cellular reprogramming (*19).* The process of reprogramming can be followed by the abundance of various cellular proteins located on the plasma membrane. During successful reprogramming, the pluripotency markers, SSEA-4 and TRA-1-60, are progressively enriched on the plasma membrane (*20).* Interestingly, SSEA-4 and TRA-1-60 appear sequentially with the latter serving as a marker of cells further along the reprogramming process and more likely to provide the rare pluripotent cells (20). Therefore, the simultaneous presence of both SSEA-4 and TRA-1-60 is an indication of cells further along in the reprogramming process (Fig. 2D, I), while cells only positive for SSEA-4 and lacking TRA-1-60 would be lagging in the process (Fig. 2D, II). Finally, cells with neither of these markers are the furthest from achieving the reprogrammed state (Fig. 2D, III) (*19*, *21*, *22*). Based on the distinction of the different reprogramming states using these makers, we analyzed the levels of HSPA5-GFP at different time points of reprogramming to ask if the UPR^ER^ induction correlated with increased reprogramming efficiency. Consistently, and robustly, we observed the highest levels of HSPA5-GFP in the cells that had progressed the furthest in the reprogramming process, the SSEA-4 and TRA-1-60 double-positive cells (Fig. 2E).

To validate the UPR^ER^ GFP reporter, we sorted the three populations (I, II, and III) at day 7 of reprogramming, a time when the UPR^ER^ is normally and transiently induced (Fig. 1C) and measured UPR^ER^ induction levels by mRNA levels of UPR^ER^ target genes (*XBP1s, HSPA5* and *GRP94).* As expected, we found the highest level of UPR^ER^ target gene induction in the SSEA-4+/TRA-1-60+ cells (I population, Fig. S3B). Additionally, we confirmed that the SSEA-4+/TRA-1-60+ population (I) was the most progressed towards reprogramming by analyzing the reactivation of endogenous pluripotency marker genes (Fig. S3C). Taken together, cells with a more advanced state of reprogramming also contained the highest induction of the UPR^ER^ by multiple measurements, indicating that proficiency of reprogramming is consistent and corollary with UPR^ER^ induction.

### Activation of the UPR^ER^ increases reprogramming efficiency

Because of the correlation between increased UPR^ER^ induction and progression towards the reprogrammed state, we asked what role, if any, did the UPR^ER^ play in the reprogramming process. To address this question, we modulated the UPR^ER^ during reprogramming either pharmacologically or genetically. Pharmacologically, we transiently activated the UPR^ER^, during periods when the UPR^ER^ is normally activated in many, but not all cells (described in detail below), using APY29, a drug that activates the RNAse activity of IRE1 (23) (Fig. S4A). Strikingly, early and transient activation of the UPR^ER^ with APY29 during the period when the UPR^ER^ is normally activated during reprogramming (days 4-7), increased the percentage of cells expressing the SSEA-4 and TRA-1-60 markers, the most mature in the reprogramming process (Fig. 3A). Importantly, to rule out that this could be due to increased rates of cell proliferation, we measured cellular proliferation in our experiments and found that it was not increased (Fig. S4B).

**Fig. 3:**
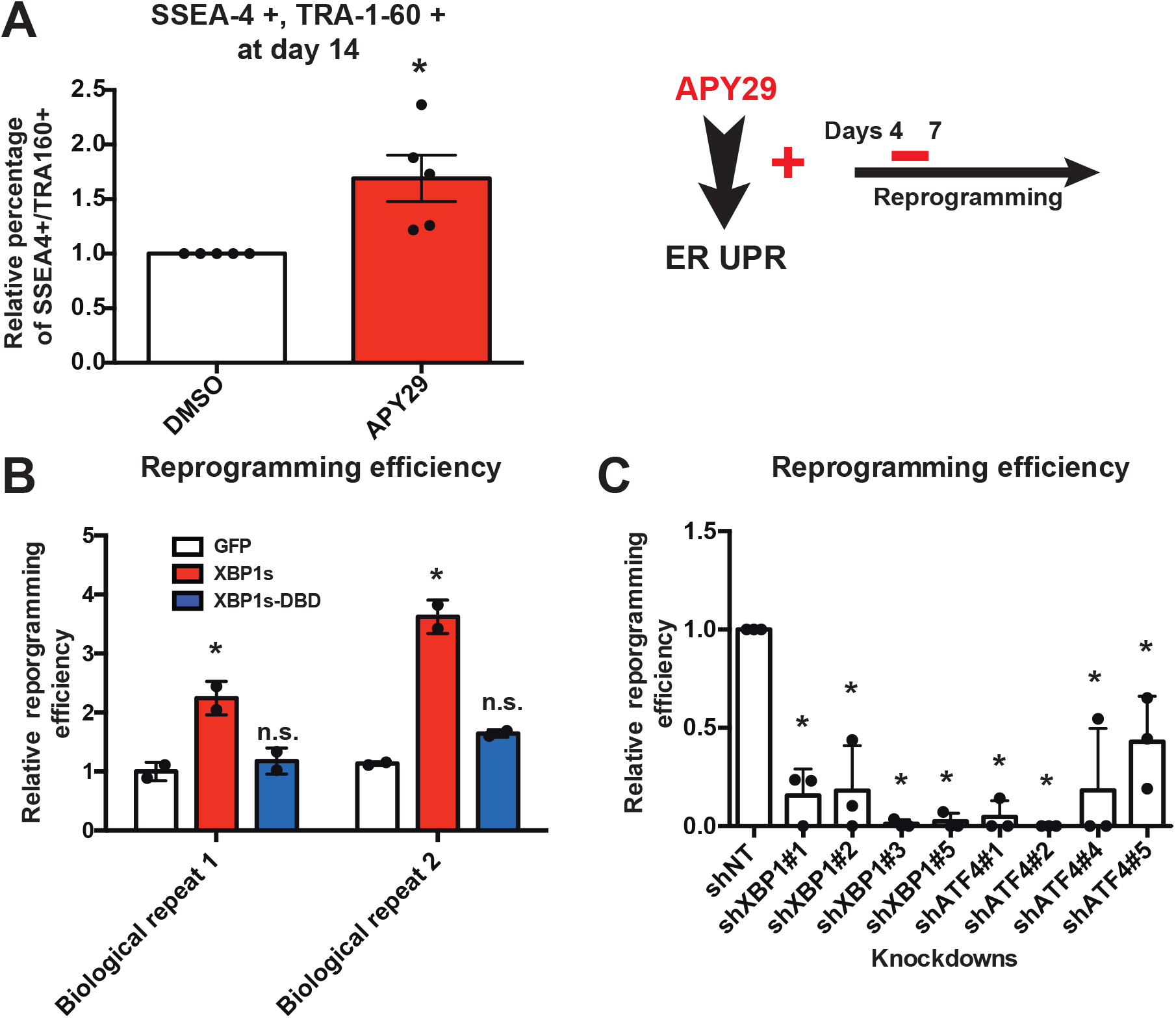
Ectopic activation of the UPR^ER^ increases the reprogramming efficiency. (A) Percentage of SSEA-4+/TRA-1-60+ cells at day 14 of reprogramming after drug treatment with APY29 (0.625 μM), an inducer of the UPR^ER^, from day 4 to day 7 of reprogramming (n=5, average +/− SEM). * indicates statistical significant difference (p-value<0.05) using an unpaired two-tailed t-test. (B) Relative reprogramming efficiency of keratinocytes measured by colony TRA-1-60 staining after 3 weeks in culture upon overexpression of emGFP, XBP1s and XBP1s-DBD (missing its DNA binding domain) with the EF1α promoter, shown are two biological replicates done in duplicate, average +/− SD. * indicates statistical difference (p-value<0.05) using a Dunnett’s multiple comparison test to the control. (C) Relative reprogramming efficiency of keratinocytes measured by colony TRA-1-60 staining after 3 weeks in culture upon knockdown of XBP1 and ATF4 (n=3, average +/− SD). * indicates statistical difference (p-value<0.05) using a Dunnett’s multiple comparison test to the control.

Intrigued by the positive and transient pharmacological manipulation of the UPR^ER^ upon reprogramming, we investigated whether genetic overexpression of XBP1s could increase cellular reprogramming efficiency. Consistent with the previous pharmacological results, overexpression of XBP1s increased reprogramming efficiency. This increased efficiency was dependent upon the transcriptional activity of XBP1s since overexpression of a mutant version of XBP1s that lacked the DNA binding domain was unable to promote reprogramming (Fig. 3B). Furthermore, we confirmed that the increased reprogramming efficiency was not caused by a higher proliferation rate due to XBP1s overexpression (Fig. S4C). Conversely and complementary, knockdown of either XBP1 or ATF4 by multiple, distinct shRNAs significantly reduced the efficiency of reprogramming (Fig. 3C, S4D and S4E). Lastly, the increased number of iPSCs created by overexpression of XBP1s were indeed pluripotent based on their ability to express pluripotency genes and differentiate into teratomas comprised of cells formed from all three germ layers as well as directly differentiate them into cells of the mesodermal and endodermal lineage (Fig. 4 and S5). We were also able to expand these observations by reprogramming primary human fibroblast using an episomal reprogramming approach (24) (Fig. S6).

**Fig. 4:**
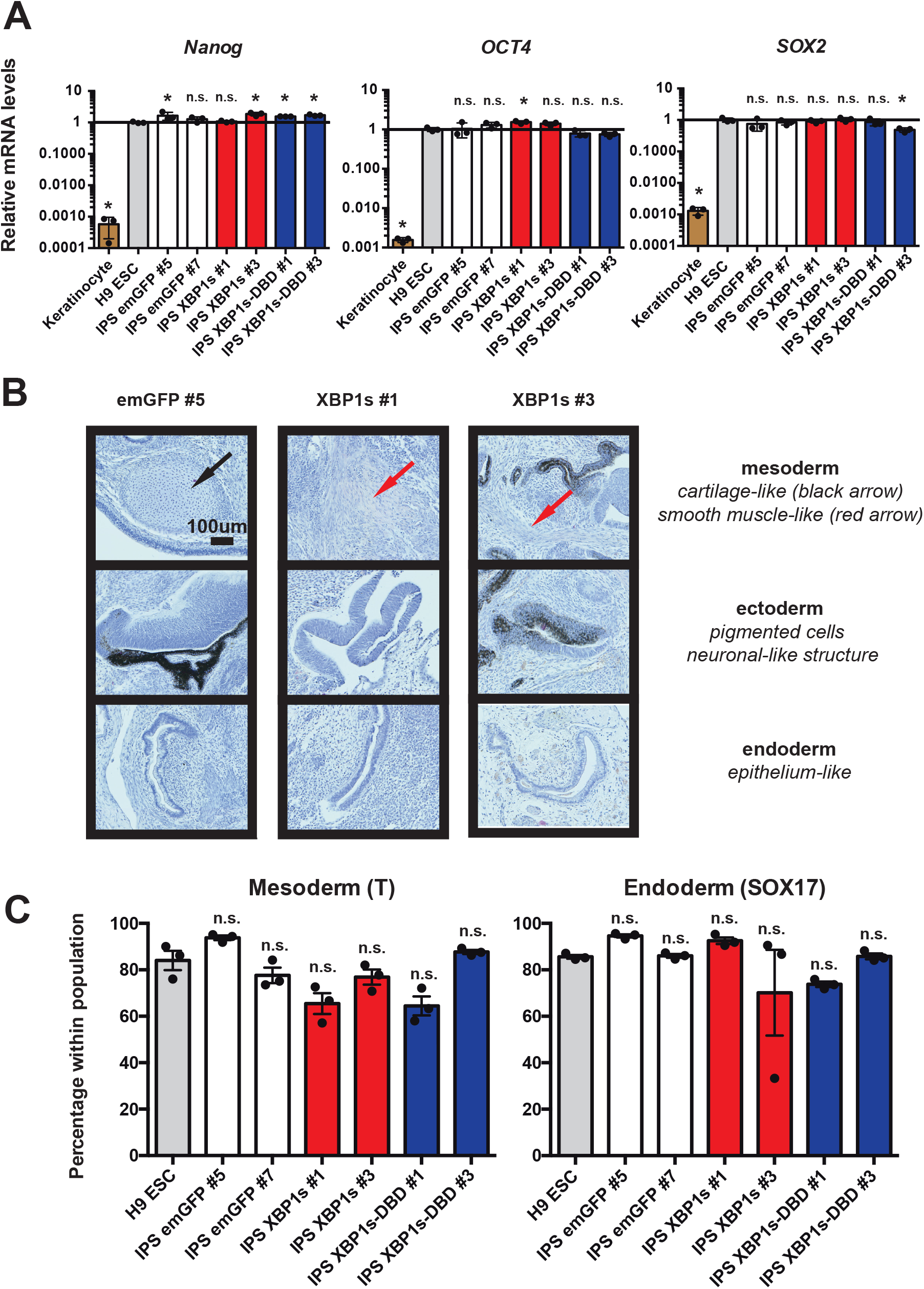
Derived iPSCs express their endogenous pluripotent genes and are pluripotent. (A) Relative endogenous mRNA levels of pluripotent genes in the derived iPSC lines relative to *GAPDH* determined by qRT-PCR (n=3, average +/− SD). Values for H9 ESCs were set to 1. * indicates statistical difference (p-value<0.05) using a Dunnett’s multiple comparison test to the control H9 ESC. (B) Hematoxylin and eosin staining of teratomas showing the three germ layers: mesoderm, ectoderm and endoderm. Teratoma formation assays were performed after confirmation of the exogenous XBP1s silencing (See Fig. 5A). (C) Directed lineage specific differentiation efficiencies assed by the percentage of cells expressing Brachyury (T) for mesoderm and Sox17 for endoderm differentiation by flow cytometry (n=3, average +/− SEM). n.s. indicates non-statistical difference (p-value<0.05) using a Dunnett’s multiple comparison test to the control H9 ESC.

Taken together, UPR^ER^ activation is not only necessary, but it is also sufficient to promote reprogramming of somatic cells to a pluripotent state. On the basis of these results we conclude that at least one arm of the proteostasis network, the UPR^ER^, plays a vital role in the cellular re-identification process of cellular reprograming.

### Activation of the UPR^ER^ during reprogramming is transient

Interestingly, after isolating iPSC colonies for characterization, we observed that the ability to properly spread and expand iPSC cells was lower when cells overexpressed XBP1s driven by the EF1α promoter with retroviral reprogramming (data not shown). These colonies remained rounded after isolation, leading to their subsequent loss in culture. On the contrary, using the episomal reprogramming method, successful iPSC clonal derivation was very similar between the GFP control and XBP1s overexpression driven by a CMV promoter (data not shown), albeit the XBP1s iPSC colonies were more numerous. The EF1α promoter is rarely silenced in embryonic stem cells, contrary to the CMV promoter (25). This led us to postulate that sustained high levels of XBP1s in iPSCs, by expression using the EF1α promoter, prevents proper spreading and expansion; and that the UPR^ER^ is required only transiently during reprogramming. Furthermore, UPR^ER^ activation could be detrimental to the fully formed iPSC, consistent with our analysis of transient UPR^ER^ activation during reprogramming (Fig. 1C, S2A and 3A). Consistent with this hypothesis, all of the EF1α promoter driven XBP1s iPSCs derived clones had similar *XBP1s* levels compared to the emGFP iPSC lines, suggesting silencing of the ectopic XBP1s transgene driven by the EF1α promoter (Fig. 5A). Additionally, the XBP1s-DBD (coding for the transcriptionally inactive XBP1s) iPSC derived clones, driven by the same EF1α promoter, did not downregulate the XBP1s-DBD transgene (Fig. 5A). XBP1s-DBD transgene does not induce the UPR^ER^. Consistent with these observations, iPSC derived clones from emGFP overexpression remained fluorescent (data not shown). The transgene was only silenced when it led to activation of the UPR^ER^ (XBP1s overexpression). Accordingly, overexpression of XBP1s using the EF1α promoter in H9 ESCs caused abnormal colony morphology with a higher density of cells within the colony with no clear colony edges and a more tridimensional growth pattern (Fig. S7A). Notably, basal levels of UPR^ER^ activity were similar, or lower, in embryonic stem cells compared to their differentiated counterparts as measured by HSPA5-GFP levels (Fig. 5B) and protein levels of ATF4, ATF6 and HSPA5 (Fig. 5C). These observations are consistent with transcriptome analyses of cellular reprogramming (26) (Table S1). Furthermore, we observed that cells undergoing cellular reprogramming transiently activated the UPR^ER^ as analyzed by both mRNA (Fig. 5D) and protein levels (Fig. 1C and S2A). Therefore, activation of the UPR^ER^ must be transient during reprogramming and appears detrimental once the cell achieves pluripotency.

**Fig. 5:**
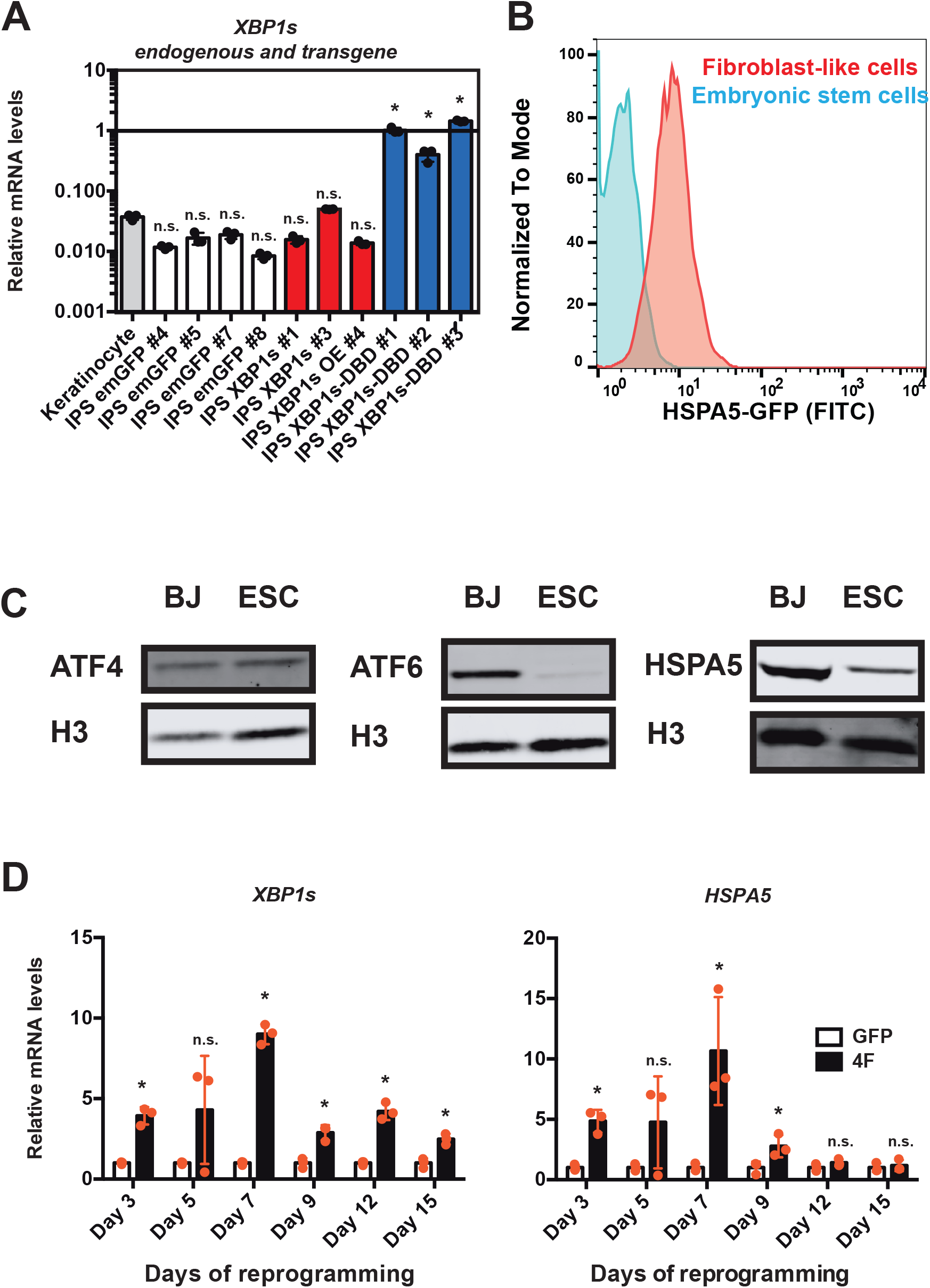
Transient activation of the UPR^ER^ is necessary during reprogramming. (A) Relative mRNA levels of *XBP1s* relative to *GAPDH* determined by qRT-PCR in iPSC colonies derived from either *emGFP, XBP1s* or *XBP1s-DBD* driven by EF1α promoter (n=3, average +/− SD). NB: this primer set will also recognize the *XBP1s-DBD* form. * indicates statistical difference (p-value<0.05) using a Dunnett’s multiple comparison test to the control keratinoytes. (B) Flow cytometry analysis of HSPA5-GFP in ESC HSPA5-GFP and the differentiated fibroblast-like cells. (C) Western blot analysis of ATF4, ATF6 and XBP1 in pluripotent stem cells and fibroblasts. Equal number of cells was loaded. (D) Relative mRNA levels of *XBP1s* and *HSPA5* relative to *GAPDH* determined by qRT-PCR during the course of cellular reprogramming (n=3, average +/− SD). GFP control was set to 1 for each day. * indicates statistical significant difference (p-value<0.05) using an unpaired two-tailed t-test.

### Levels of UPR^ER^ activation positively correlate with the reprogramming efficiency

Because reprogramming efficiency could be increased by the activation of the UPR^ER^ and decreased by the loss of XBP1 or ATF4, we postulated that the ability to ectopically induce the UPR^ER^ of individual, genetically identical, somatic cells cultured in identical conditions could be stochastic and might also outline part of the variable nature of the process of cellular reprogramming. To address this question, we followed the induction of the HSPA5-GFP reporter within individual cells in a population undergoing cellular reprogramming. We found a Gaussian distribution of HSPA5-GFP fluorescence amongst the cell population undergoing reprogramming (Fig. 6A), indicating that UPR^ER^ activation was variable across the cell population. To test whether the intrinsic ability of a cell to induce the UPR^ER^ was predictive of further success along the reprogramming process, we subdivided the Gaussian distributed HSPA5-GFP population at day 8 of reprogramming into 3 equal subpopulations (low, medium and high UPR^ER^ induction via

**Fig. 6:**
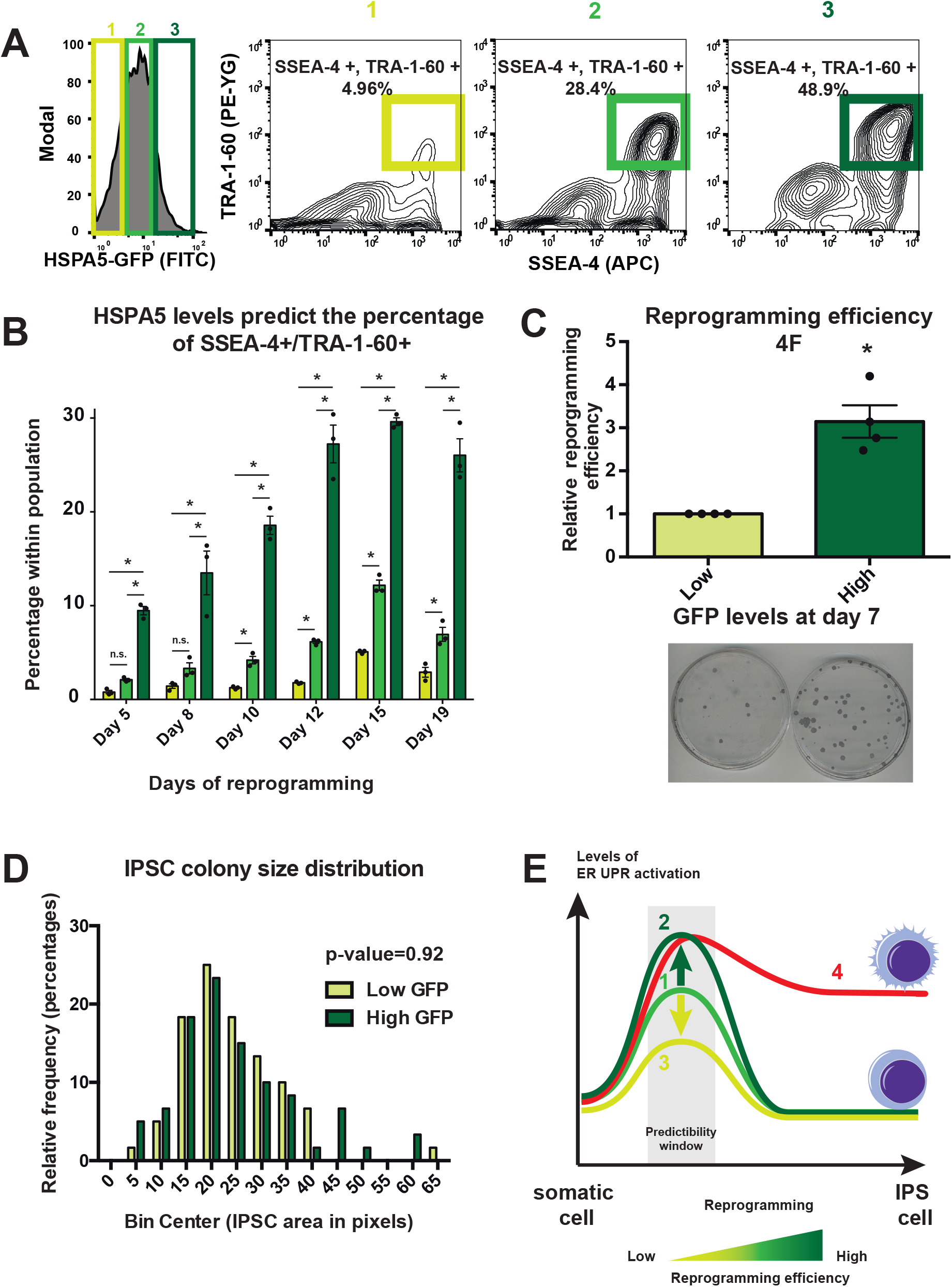
HSPA5-GFP levels predict the reprogramming efficiency. (A) Histogram of fibroblast-like HSPA5-GFP at day 8 of reprogramming. 1, 2, 3 subdivide the population into 3 equal parts. Each of them is represented in the right panel by their SSEA-4 and TRA-1-60 staining. The percentage of double positive cells within each of these populations is shown. (B) Percentage of SSEA-4+/TRA-1-60+ cells within each population 1, 2, 3 during reprogramming (n=3, average +/− SD). * indicates statistical difference (p-value<0.05) using Newman-Keuls multiple comparison test between all the conditions for each day. (C) Upper panel shows relative reprogramming efficiency of fibroblast-like HSPA5-GFP sorted at day 7 of reprogramming based on their GFP levels and assessed by TRA-1-60 colony staining (n=4, average +/− SEM). Lower panel shows a representative picture of the staining. * indicates statistical difference (p-value<0.05) using an unpaired two-tailed t-test. (D) iPSC colony size distribution from experiment Fig. 6C. The area of ~60 iPSC colonies was measured with ImageJ. There was no significant difference in the mean colony size (unpaired two-tailed t-test). (E) Cells undergoing a successful cellular reprogramming activate the UPR^ER^ transiently (1). Increased levels of UPR^ER^ activation increase the efficiency of cellular reprogramming (2) while decreasing the UPR^ER^ activation negatively impacts the efficiency of cellular reprogramming (3). In our experimental design, it is possible to predict the efficiency of reprogramming based on the levels of UPR^ER^ activation around day 7, depicted by the predictability window. Cells unable to decrease the levels of UPR^ER^ activation give rise to iPSCs unable to properly spread (4).

GFP levels) (Fig. 6A). We found that the percentage of cells most progressed and more likely to form iPSCs, SSEA-4+/TRA-1-60+ cells, was highest in cells with the highest levels of HSPA5-GFP (Fig. 6A). We expanded this observation to multiple time points during reprogramming and found that we could not break the correlation between UPR^ER^ induction and increased reprogramming efficiency (Fig. 6B).

To further interrogate the predictive value of the UPR^ER^ induction and iPSCs formation, we sorted cells at day 7 of reprogramming based on their levels of HSPA5-GFP into two populations, high and low levels (Fig. S7B), and assessed iPSC colony formation. After 10 days in culture, cell colonies were stained for TRA-1-60. As predicted, cells with higher levels of HSPA5-GFP at day 7 gave rise to more iPSC colonies (Fig. 6C). We next tested if the reprogramming process was faster in cells with higher levels of UPR^ER^ induction. We reasoned that if this was the case, then the size of the colonies would be larger compared to cells with low UPR^ER^ induction. We measured the area of the iPSC colonies from (Fig. 6C) and there were no significant differences in size of the colonies, suggesting that cells with higher UPR^ER^ induction do not reprogram faster (Fig. 6D). These finding indicate that the intrinsic ability of a somatic cell to induce the UPR^ER^ is predictive of its likelihood of becoming pluripotent.

Because c-MYC is a proto-oncogene that facilitates genomic instability and its ectopic overexpression could lead to deleterious side effects during transplantation of iPSCs into hosts, we asked if we could bypass the need for c-MYC using the intrinsic induction of the UPR^ER^ in combination with three of the four reprogramming factors, OCT4, SOX2 and KLF4 (3F). We found high levels of UPR^ER^ induction in cells that had progressed the furthest in the reprogramming process using only 3 factors (SSEA-4/TRA-1-60+), consistent with our analysis with all four factors (Fig. S8A). Additionally, higher HSPA5-GFP correlated with increased percentage of SSEA-4/TRA-1-60+ cells (Fig. S8B). Consistent with this observation, cells with higher levels of HSPA5-GFP at day 7 gave rise to more iPSC colonies (Fig. S8C), indicating that induction of the UPR^ER^ could be used as an alternative approach to circumvent potential off-target side effects that might be negative when creating iPSCs with potential pro-oncogenes.

## Discussion

The details and mechanics required for successful cellular reprogramming are becoming more apparent, but the low efficiency due to its stochastic nature are less well understood (*3*). The extremely low efficiency can be augmented by the addition of supplementary factors such as other pluripotency-associated factors, cell cycle-regulating genes and epigenetic modifiers (4). However, none of these factors address the role of the proteostasis network or organelle integrity as an important driver for reprogramming. We find that an early ER stress is an essential step for a somatic cell to reprogram and that ectopic, transient activation of the UPR^ER^ increases reprogramming efficiency. Moreover, the stochastic nature of the reprogramming process could be partially explained by the ability of a cell to properly mount an ER stress response and can be used as a predictive marker of successful reprogramming (Fig. 6E).

We were surprised that XBP1s can robustly increase the reprogramming efficiency and its requirement was not previously observed in reprogramming paradigms. One explanation could be that XBP1s is transiently upregulated during reprogramming. Additionally, XBP1 activation requires a regulatory splicing event, while most of the other reprogramming factors were inferred based on their high levels in the ESCs. Indeed, transient activation of the UPR^ER^ during the early phase of reprogramming using the IRE1 activating drug, APY29, was sufficient to increase its efficiency.

Interestingly, XBP1s, among other UPR^ER^ effectors, is required during development and differentiation (14). Thus, consistent with the original hypothesis to identify reprogramming factors utilizing genes required for normal development and differentiation could help reprogram better by enabling a successful transition between the two cell states. Therefore, it is intriguing that a central player of the proteostasis network plays an essential role not only in development, but also reprogramming.

The mechanism through which the activation of the UPR^ER^ increases reprogramming efficiency remains to be elucidated. The UPR^ER^ activation leads to a global reduction of protein synthesis (*10*) and the degradation of mRNA associated with the ER membrane (27). One possibility is that the ER proteome is cleared from a substantial part of its somatic signature allowing the new pluripotent proteome to be set. Therefore, the activation of the UPR^ER^ must be transient, which is supported by our findings. Additionally, the ectopic activation of the UPR^ER^ could provide a cytoprotective buffer to explore different states and consequently reach pluripotency without inducing apoptosis during the reprogramming process. Consistent with this idea, cells with a higher level of UPR^ER^ activation reprogram at a higher rate. Interestingly, although less explored, bidirectional regulation between DNA-damage responses and the UPR^ER^ have been shown (*28).* A cluster of DNA damage and DNA repair genes was identified as a direct target of XBP1s (*29).* Although not explored here, this could allow the potential to counter the negative effects of c-MYC, a well-known inducer of genomic instability, without affecting negatively the reprogramming efficiency as supported by our data. While 3-Factor reprogramming had increased efficiency when the UPR^ER^ was activated, we question that replacing c-MYC with XBP1s would meet the needs of the field, as 3-Factor reprogramming is extremely inefficient. Instead, the combination of a transient drug to induce the UPR^ER^, such as APY29, in combination with the four Yamanaka factors may prove extremely useful for clinical applications and cells difficult to reprogram.

We predict that effectors ensuring protein quality control can be potent facilitators of reprogramming in assisting the transition from one cell state to another. Previous work in our lab (30) and others (31) has already linked protein quality control through the ubiquitin-proteasome system with stem cell maintenance, differentiation and reprogramming. The role of other regulatory elements of protein quality control such as the mitochondrial unfolded protein response (UPR^mt^), and molecular chaperones involved in the heat shock response remain largely unexplored in the regulation of stem cell differentiation or reprogramming. How these processes are involved in reprogramming, as well as their potential cross-play with the UPR^ER^, will need to be explored. We believe that our observations could also be relevant to transdifferentiation paradigms.

## Material and methods

### Cell culture

Human dermal fibroblasts (Lonza CC-2511 and CC-2509), HEK293FT (ThermoFisher, R70007), BJ human fibroblasts (ATCC, CRL-2522), fibroblast-like cells and irradiated CF-1 mouse embryonic fibroblasts (GlobalStem) were grown in DMEM, 10% FBS, 1x Pen/Strep, 1x glutamax and 1X non-essential amino acids (NEAA) (all from Invitrogen).

The hESC line H9 (WA09, WiCell Research Institute) and the other hiPS generated lines were cultured with mTeSR1 media (STEMCELL™ Technologies) on Geltrex (Invitrogen). Human keratinocytes (Lonza 192907) were cultured with KGM-Gold media (Lonza).

### Plasmids

A list of the plasmids and the cloning strategy can be found in Table S2.

### Viral production

Lentiviral and moloney-based retroviral pMX-derived vectors were co-transfected with their respective packaging vectors in 293FT cells using JetPrime transfection reagent to generate viral particles as previously described (*18*). The viral supernatant was filtered through a 0.45 μM filter.

### iPSC generation

Primary cells were spinfected with the viral supernatant containing the reprogramming factors and other factors during 1hour at 1000g in presence of 8μg/mL of polybrene (Millipore) twice, 24 hours apart. The regular media was replaced after each round. Selection was started the next day of the last transfection, 48 hours later cells were dissociated with TrypLE (Invitrogen) and plated on top of irradiated MEFs in their regular media. The next day cells were switched to iPS media containing DMEM/F12, 20% knockout serum replacement, 1X Pen/Strep, 1X glutamax, 1X NEAA, 10ng/mL bFGF (all from Invitrogen), and 55μM β-mercaptoethanol (Sigma). To evaluate the reprogramming efficiency, the number of plated cells was counted, after 2-3 weeks cells were fixed with 4% PFA and stained for TRA-1-60 as previously described (32) and scored. Briefly, fixed cells were blocked for 1 hour at room temperature in 1xPBS, 3% FBS, 0.3% Triton X-100, then incubated with biotin-anti-TRA-1-60 (eBioscience13-8863-82, 1:250) over night at 4C and the next day streptavidin horseradish peroxidase (Biolegend 405210, 1:500) for 2 hours at room temperature. Staining was developed with the sigmaFast DAB kit (D0426). Alternatively, an alkaline phosphatase (AP) staining was performed for episomal reprogramming experiments as instructed by the Millipore detection kit (SCR004). Briefly, cells were fixed in 4% PFA for less than a minute to avoid losing the AP activity. Cells were rinsed with TBS-T and covered with Fast Red Violet Solution/water/Naphthol (2:1:1) for 20 min followed by a wash with PBS. AP positive colonies were then counted.

For time course studies, imaging and flow cytometry, cells were plated on geltrex coated plates instead of MEFs.

Where indicated, after plating on geltrex, cells were incubated with APY29 (Chem Scene, CS-2552) for 3 days.

Alternatively, cells were also reprogrammed using an episomal electroporation system (24). Briefly, cells were first selected with the appropriate factor. 500,000 cells were then electroporated with the episomal constructs using the nucleofector kit (Lonza, VPD-1001). Cells were plated and kept in their original media. After 6 days, cells were dissociated and plated on freshly plated MEFs. Cells were switched to iPS media the next day.

### Derivation of fibroblast-like cells

Stem cells were differentiated into fibroblast-like cells using an embryoid body (EB)-mediated protocol. Stem cells grown on Geltrex were detached using dispase, resuspended in DMEM/F12, 20% FBS, 1x glutamax, 1x NEAA, 1x Pen/Strep and 55μM β-mercaptoethanol and grown on low adhesion plates for 4 days with media change. EBs were plated on gelatin-coated plates and cultured with the same media. When EBs spread and cells appeared fibroblast looking, the culture was dissociated using TrypLE and replated using a regular fibroblast media. This was serially done until the whole population became uniform.

### RNA isolation and real-time PCR

Cells were collected in Trizol^®^. A classic chloroform extraction followed by a 70% ethanol precipitation was performed. The mixture was then processed through column using the RNeasy quiagen kit as described by the manufacturer. Quantitect reverse transcription kit (Quiagen) was used to synthesize complementary DNA. Real-time PCR was performed using Sybr select mix (Life Technologies). *GAPDH* expression was used to normalize gene expression values. Primer sequences can be found in the Table S2.

### Western blot analysis

Cells were washed with PBS and RIPA buffer was added to the plates on ice. Cells were scraped, collected and stored at −20C. The RIPA buffer was always supplemented with Roche cOmplete mini, and phosSTOP when needed. 20 μg of protein was loaded per lane and actin or histone H3 were used as a loading control in precast 4-12% Bis-Tris NuPage gels (Invitrogen). Proteins were blotted on nitrocellulose membranes using the NuPage reagents according to the manufacturer instructions. Membranes were prepared for imaging using Odyssey^®^ CLx Imaging System-LI-COR Biosciences with the appropriate reagents. Briefly, membranes were incubated in the proprietary blocking buffer for 1 hour at room temperature. Overnight primary antibody incubation at 4C was done using the blocking buffer and 0.1% Tween-20. Membranes were washed in TBS-T then incubated with secondary antibody for 1 hour at room temperature. Membranes were then washed in TBS-T with a final wash in TBS. The software ImageStudio was used to quantify the band intensities. For the list of antibodies and concentrations refer to Table S2.

### Fluorescent immunostaining

Cells on slides were fixed with 4% PFA for 15min and washed with PBS. 2% donkey-serum blocking buffer in PBS was used for 1 hour at room temperature. Primary antibody incubation was done overnight. After PBS washes, secondary antibody was added for 1 hour at room temperature. After PBS washes, slides were mounted with mounting media containing DAPI. For the list of antibodies and concentrations refer to Table S2.

### Flow cytometry

For cell analysis, cells were dissociated with TrypLE and pelleted. 100 μL of a fluorescent-conjugated antibodies cocktail (5 μL of SSEA-4 330408, 5 μL of TRA-1-60 330610 Biolegend) in staining media (1xPBS, 2% FBS) was used to resuspend the pellet and incubated 30min on ice. Cells were then resuspended in excess of staining media, span down and resuspended in staining media, filtered through a cell strainer and kept on ice. Cells were analyzed using the BD Bioscience LSR Fortessa. The analysis was done using the FlowJo software. For directed mesodermal and endodermal differentiation experiments a similar workflow was used with the exceptions of using accutase to dissociate the cells and using saponin buffer (1mg/mL saponinin PBS +1% BSA) to permeabilize the cells before incubating with fluorescent-conjugated SOX17 (BD biosciences 562594) or Brachyury (Fisher Scientific IC2085P) in saponin buffer.

For cell sorting, a similar procedure was followed. Cells were eventually resupsended in their media supplemented with rock inhibitor and sorted accordingly using the BD Bioscience Influx Sorter (Fig. S7B). Cells were then transferred to appropriate dishes for culture and kept on rock inhibitor during the next 24 hours.

### ER secretion assay

Transduced cells with Gluc-CFP were incubated 24 hours with fresh media and the supernatant was collected for analysis. An equal volume of Gluc Glow buffer (nanolight) was added to the supernatant in a 96-well plate format. The luminescence was measured by a TECAN plate reader and integrated over 50 ms.

### Cell assay

Cells were plated on 96-well plates and treated with the appropriate condition. After the desired incubation time, cell titer glow buffer (Promega) was added to the wells (1:5 volume) and incubated for 12min on a shaker. The luminescence was measured with the TECAN plate reader and integrated over 1s.

### Electron microscopy

Cells were fixed with 2% glutaraldehyde in 0.1 M phosphate buffer for 5 min. Samples were rinsed with 0.1M sodium cacodylate Buffer (3×5 min) followed by the addition of 1% osmium tet, 1.5% ferrocyanide in 0.1M cacodylate buffer (5min). After washing with water (3×5min), 2% uranyl acetate was added for 5min followed by a water rinse. A dehydration series of ethanol was then completed: 35%, 50% 75%, 100%, 100% (5 min each). A 1:1 ethanol/resin (3×10min) incubation followed by 100% resin (3×10min) was done. The samples were cured over 48hrs, sectioned at 50nm with a microtome using a Diatome. Sections were placed on a coated copper mesh grid. They were then stained with uranyl acetate for 5 min, and then stained with lead citrate for 5 min before imaging.

### Genome editing and southern blot analysis

Transcription activator-like effector nuclease (TALENs) technology was used to create a fusion HSPA5-GFP by insertion of eGFP-PGK-Puro at the 3’ end of the HSPA5 locus. We followed the protocol described in (33). TALENs were cloned to bind ACAGCAGAAAAAGATGA and ATTACAGCACTAGCA sequences and generate a double-stranded break proximal to the STOP codon. The donor plasmid OCT4-eGFP-PGK-Puro, published in (33), was adapted to target HSPA5 by changing the homology arms. H9 cells were electroporated and clonal expansion after puromycin selection was done. Successful targeting was confirmed by southern blot using the GFP probe published in (*33*). Further information can be found in Fig. S3A.

### Teratoma assay and directed differentiation

Teratoma formation assays where performed as previously described in (*21*). For directed endoderm and mesoderm differentiation we used STEMdiff™ kits from STEMCELL™ technologies and followed their instructions (CAT#05232 and 05233).

### Statistical analysis

The software Excel and Prism were used to perform the statistical tests. The corresponding statistical tests and the number of biological repeats, denoted as n, are indicated in the figure legends. When comparing only 2 conditions we used a t-test. If multiple comparisons were done, we corrected for the multiple comparisons. For example if all the conditions were compared to the control only and no other comparisons between the conditions was intended (e.g. A with B, A with C) then a Dunnett’s multiple comparison test was used. If all the conditions were compared to each other (e.g. A with B, B with C and A with C) then a Newman-Keuls multiple comparison test was used. SD and SEM stand respectively for standard deviation and standard error of the mean. For drug dose response assays, a log(drug) vs normalized response with viable slope model was used to determine the EC50.

## Acknowledgements

Funding was provided by HHMI, CIRM (RB5-06974), NIA (R01AG042679) and NIH (R01CA196884) grants.

We thank Dr B. Tannous who generously provided the Gluc-CFP, the Tjian lab for the episomal reprogramming vectors, Dr A. Panopoulos and Dr S. Ruiz for the pMX reprogramming vectors and the reprogramming protocols, R. Forster, J. Boyle, K. Hennick, B. Webster and G. Qing for technical help.

## Supplementary Materials

**Fig. S1:**
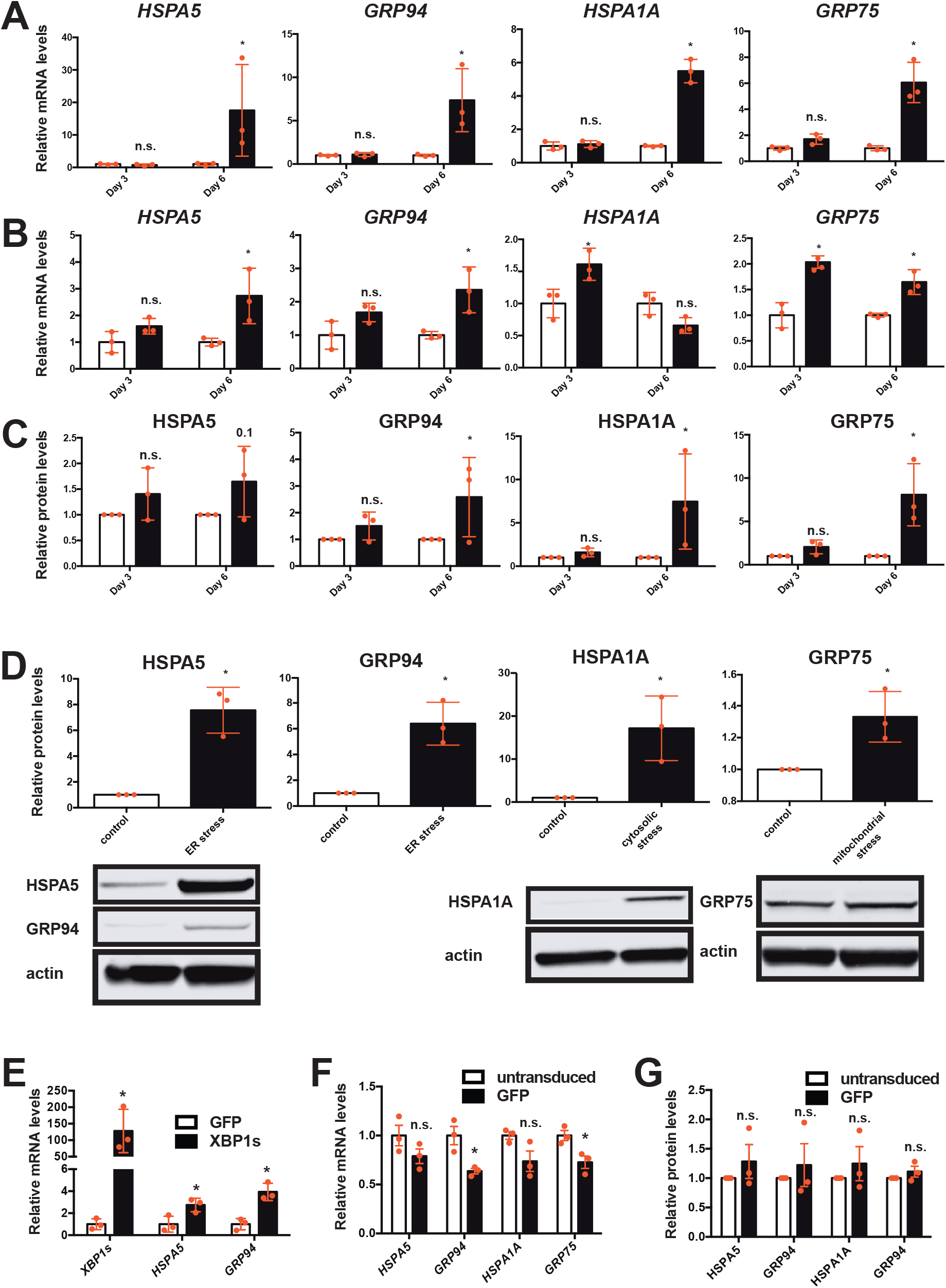
The reprogramming factors activate the three major unfolded protein responses during reprogramming. (A) Relative mRNA levels of the main effectors of the UPR^ER^ (HSPA5 and GRP94), HSR (HSPA1A) and UPR^mt^ (GRP75) relative to *GAPDH* determined by qRT-PCR (n=3, average +/− SD) during reprogramming in neonatal keratinocytes. GFP control was set to 1 for each day. * indicates statistical difference (p-value<0.05) using an unpaired twotailed t-test. (B) Relative mRNA levels of the main effectors of the UPR^ER^ (HSPA5 and GRP94), HSR (HSPA1A) and UPR^mt^ (GRP75) relative to *GAPDH* determined by qRT-PCR (n=3, average +/− SD) during episomal reprogramming with electroporation. GFP control was set to 1 for each day. * indicates statistical difference (p-value<0.05) using an unpaired two-tailed t-test. (C) Western blot band intensities quantified and normalized to actin during reprogramming from Fig. 1B. GFP control was set to 1 for each day (n=3, average +/− SD), Fisher LSD test. (D) Western blot band intensities quantified and normalized to actin after activation of the UPR^ER^ (ER stress: 0.1μg/mL tunicamycin treatment during 24h), the HSR (cytosolic stress: 42°C heat-shock for 30min with a 3 hour recovery) and UPR^mt^ (mitochondrial stress: over-expression of polyglutamine huntingtin (*34*)) (n=3, average +/− SD). Below graphs are representative western blots. * indicates statistical difference (p-value<0.05) using an unpaired two-tailed t-test. (E) mRNA levels of *XBP1s, HSPA5* and *GRP94* upon overexpression of XBP1s (n=3, average +/− SD). GFP control was set to 1. * indicates statistical difference (p-value<0.05) using an unpaired two-tailed t-test. (F) mRNA levels of *HSPA5, GRP94, HSPA1A* and *GRP75* between untransduced and GFP transduced keratinocytes (n=3, average +/− SD). * indicates statistical difference (p-value<0.05) using an unpaired two-tailed t-test. (G) Western blot band intensities quantified and normalized to actin between untransduced and GFP transduced fibroblasts from Fig. 1B (n=3, average +/− SD). n.s. indicates non statistical difference (p-value<0.05) using an unpaired two-tailed t-test.

**Fig. S2:**
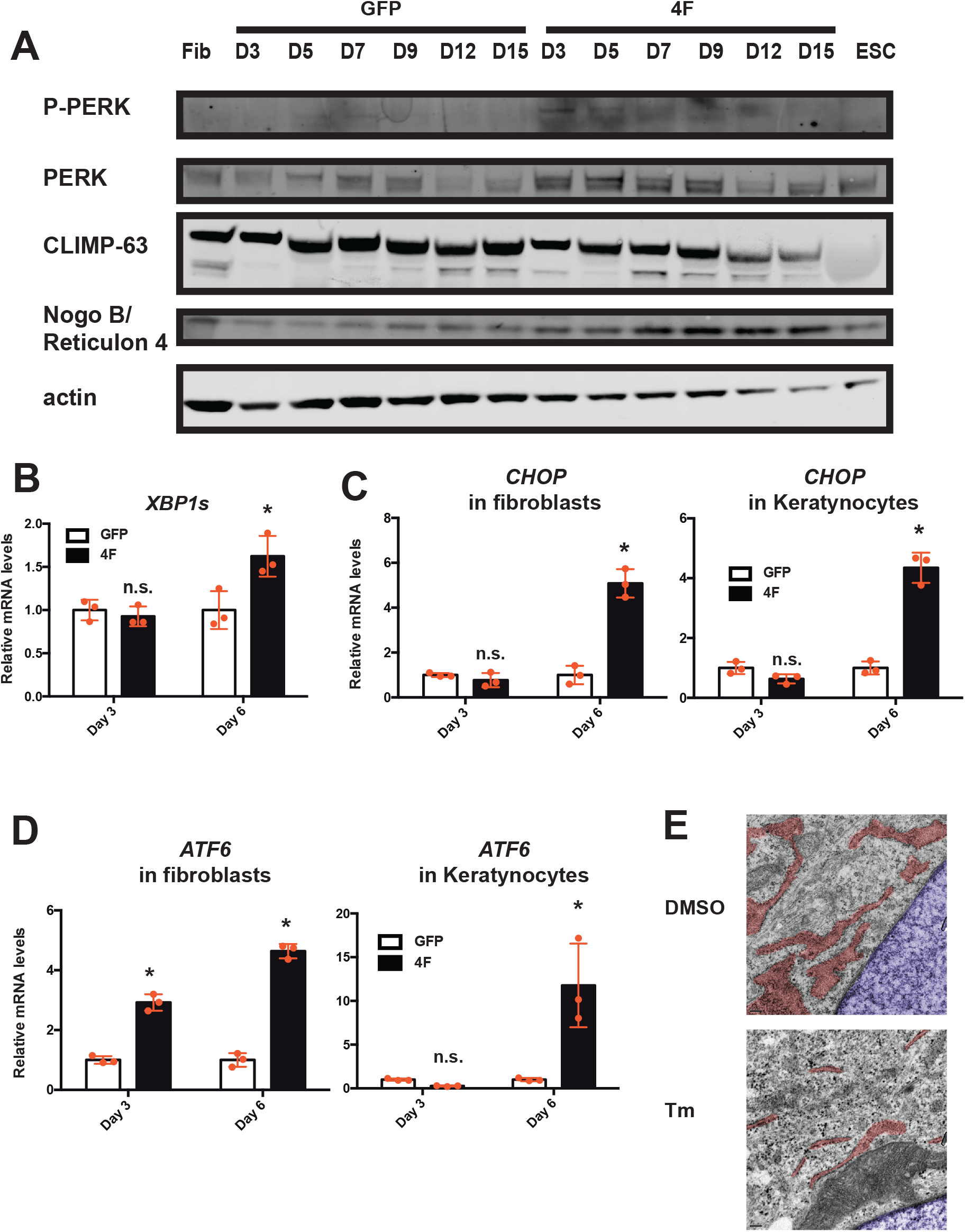
The reprogramming factors activate all the three branches of the UPR^ER^ during reprogramming. (A) Time course reprogramming western blot analysis of PERK, P-PERK, CLIMP-63, Reticulon 4 (isoform Nogo B) and loading controls. (B) mRNA levels of *XBP1s* during cellular reprogramming in neonatal keratinocytes (n=3, average +/− SD). GFP control was set to 1 for each day. (C) mRNA levels of *CHOP* a downstream target of the PERK pathway during cellular reprogramming in two different cell types (n=3, average +/− SD). GFP control was set to 1 for each day. (D) mRNA levels of *ATF6* during cellular reprogramming in two different cell types (n=3, average +/− SD). GFP control was set to 1 for each day. x(E) Electron microscopy of fibroblasts treated with 1μg/mL of tunicamycin during 24h (Tm), an ER stress inducer, scale bar = 0.2 μm. Pseudo-colors blue and red mark respectively the nucleus and the ER. * indicates statistical difference (p-value<0.05) using an unpaired two-tailed t-test.

**Fig. S3:**
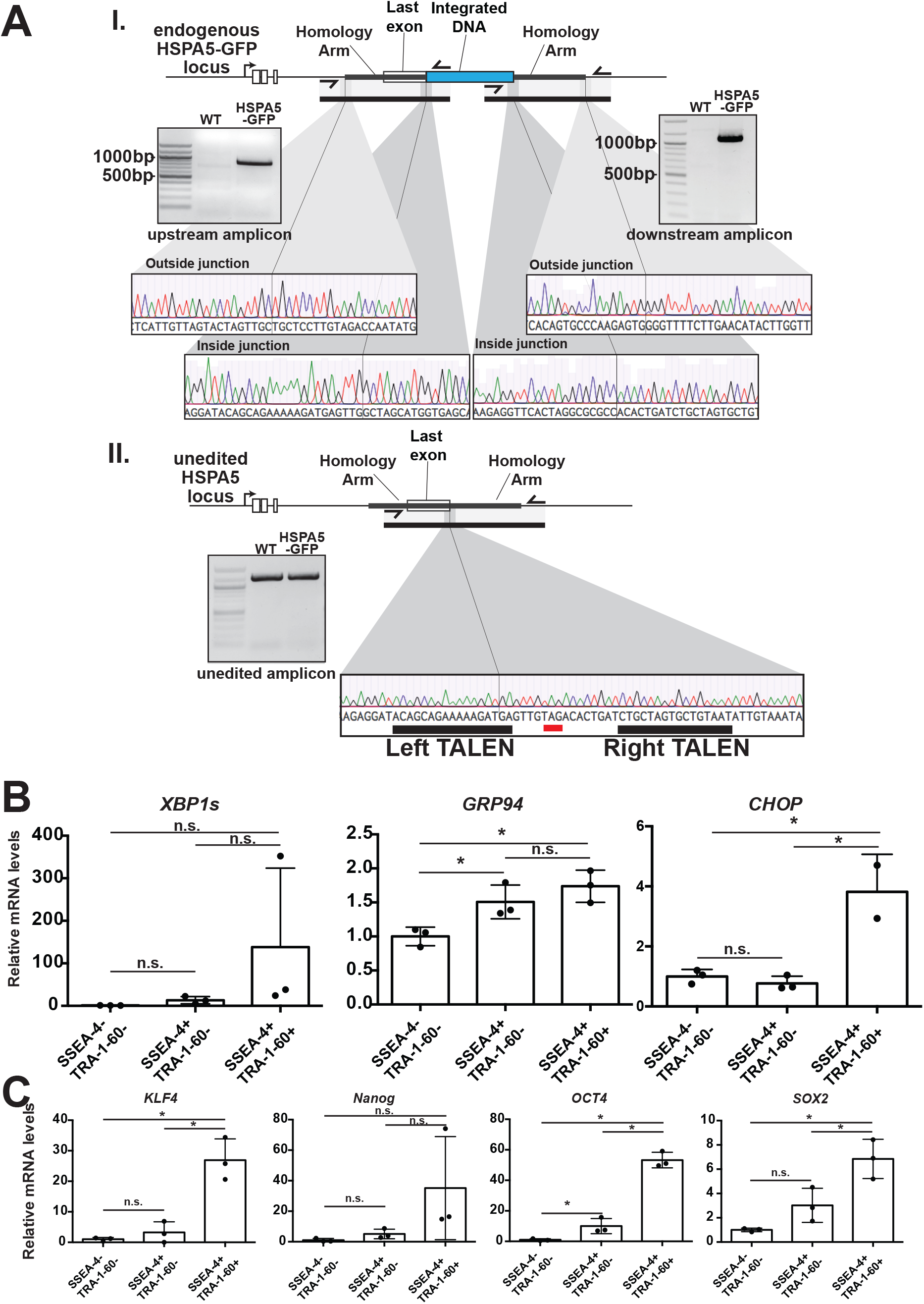
Activation of the UPR^ER^ and reactivation of the endogenous pluripotent genes during the different cellular reprogramming stages using fibroblast-like HSPA5-GFP cells. (A) I. Schematic of edited HSPA5 locus with upstream and downstream primer pairs indicated (top). One primer binds inside the integrated DNA and the opposite primer binds outside the homology arm within the HSPA5 locus. Gel electrophoresis shows that these primers amplify DNA of the expected size from the HSPA5-GFP clone only. Expected sizes: upstream (848bp), downstream (1103bp). Sequencing of the PCR product reveals the sequence expected from integration via homology dependent repair; Sanger sequencing chromatographs of the inside and outside junctions are shown (bottom). II. Schematic of unedited HSPA5 locus with primer pair indicated (top). This primer pair yields a band of the expected size and sequence when WT or HSPA5-GFP genomic DNA are used as template. The Sanger sequencing chromatograph of the HSPA5-GFP sequencing reaction at the TALEN binding sites shows WT sequence (bottom). Ladder bands from top to bottom (in bp): 100, 200, 300, 400, 500/517, 600, 700, 800, 900, 1000, 1200, and 1517. Relative endogenous mRNA levels of UPR^ER^ (B) and pluripotent genes (C) in the differentially reprogrammed populations relative to *GAPDH* determined by qRT-PCR (n=3, average +/− SD). Values for SSEA-4-/TRA-1-60-were set to 1. * indicates statistical difference (p-value<0.05) using Newman-Keuls multiple comparison test between all the conditions for each day.

**Fig. S4:**
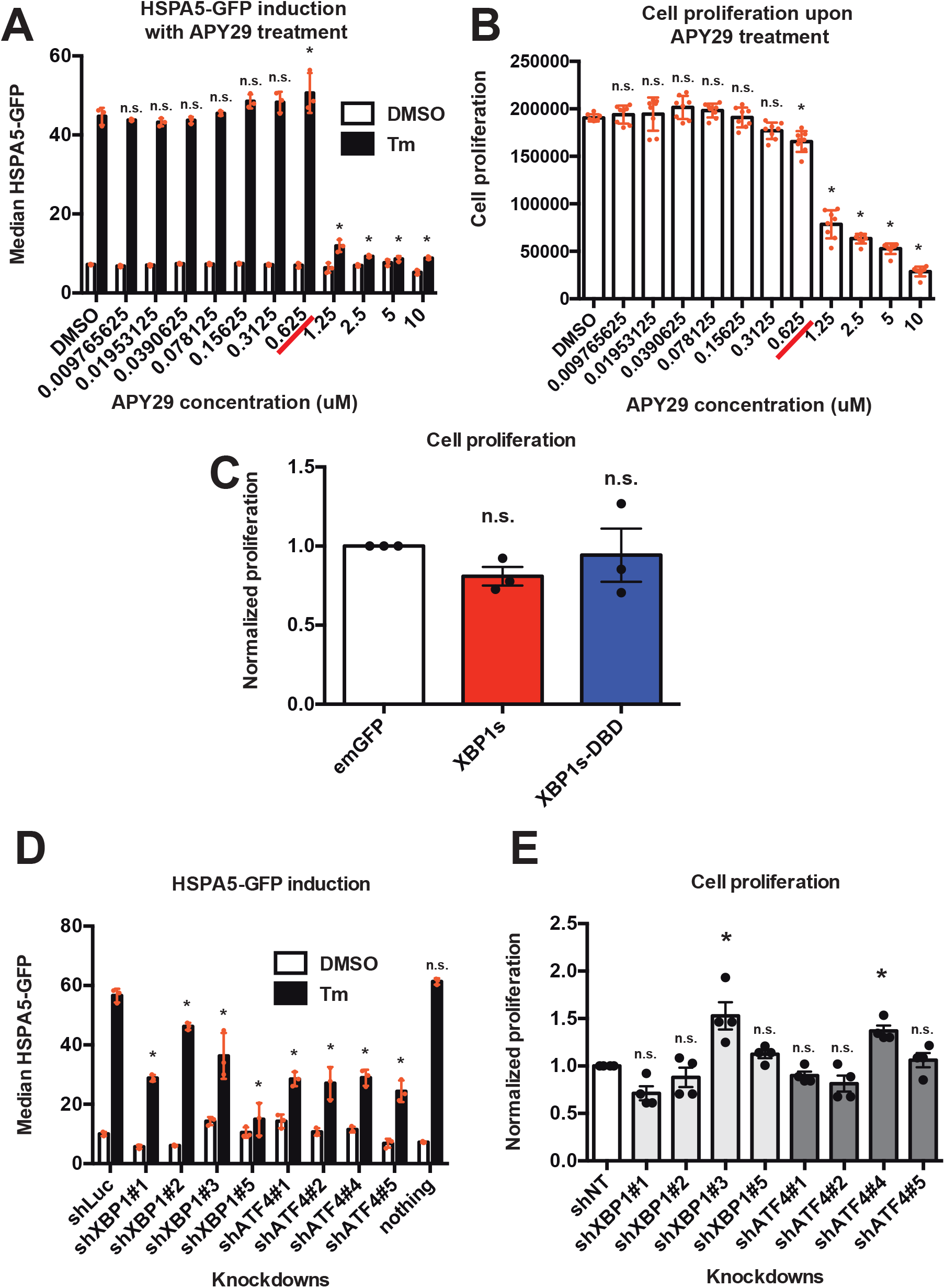
Modulation of the UPR^ER^ and its impact on cell proliferation. (A) Median HSPA-GFP levels with and without 1μg/mL tunicamycin treatment during 48 hours pretreated during 24 hours with different concentration of APY29. The drugs were kept during the entire experiment (n=4, average +/− SD). * indicates significant statistical difference (p-value<0.05) using a Dunnett’s comparison test to control DMSO Tm. (B) Growth tested by cell-titer glow assay with different concentrations of APY29 treated during 3 days (n=8, average +/− SD). The red line corresponds to the concentration used for the experiment in Fig. 2B. Error bars indicate the standard deviation. * indicates significant statistical difference (p-value<0.05) using a Dunnett’s comparison test to control DMSO. (C) Growth tested by cell-titer glow assay on keratinocytes upon expression of the 4 reprogramming factors and the overexpression of emGFP, XBP1s and XBP1s-DBD with the EF1α promoter at 3 days of reprogramming (n=3, average +/− SEM). n.s. indicates non-significant statistical difference (p-value<0.05) using a Dunnett’s comparison test to control emGFP. (D) Induction of HSPA5-GFP reporter by tunicamycin upon knockdown of either XBP1 or ATF4 analysed by flow cytometry (n=4, average +/− SD). * indicates significant statistical difference (p-value<0.05) using Dunnett’s comparison test to control shLuc. (E) Growth tested by cell-titer glow assay on keratinocytes upon knockdown of either XBP1 or ATF4 after 3 days of culture during reprogramming (n=4, average +/− SEM). * indicates significant statistical difference (p-value<0.05) using a Dunnett’s comparison test to control shNT.

**Fig. S5:**
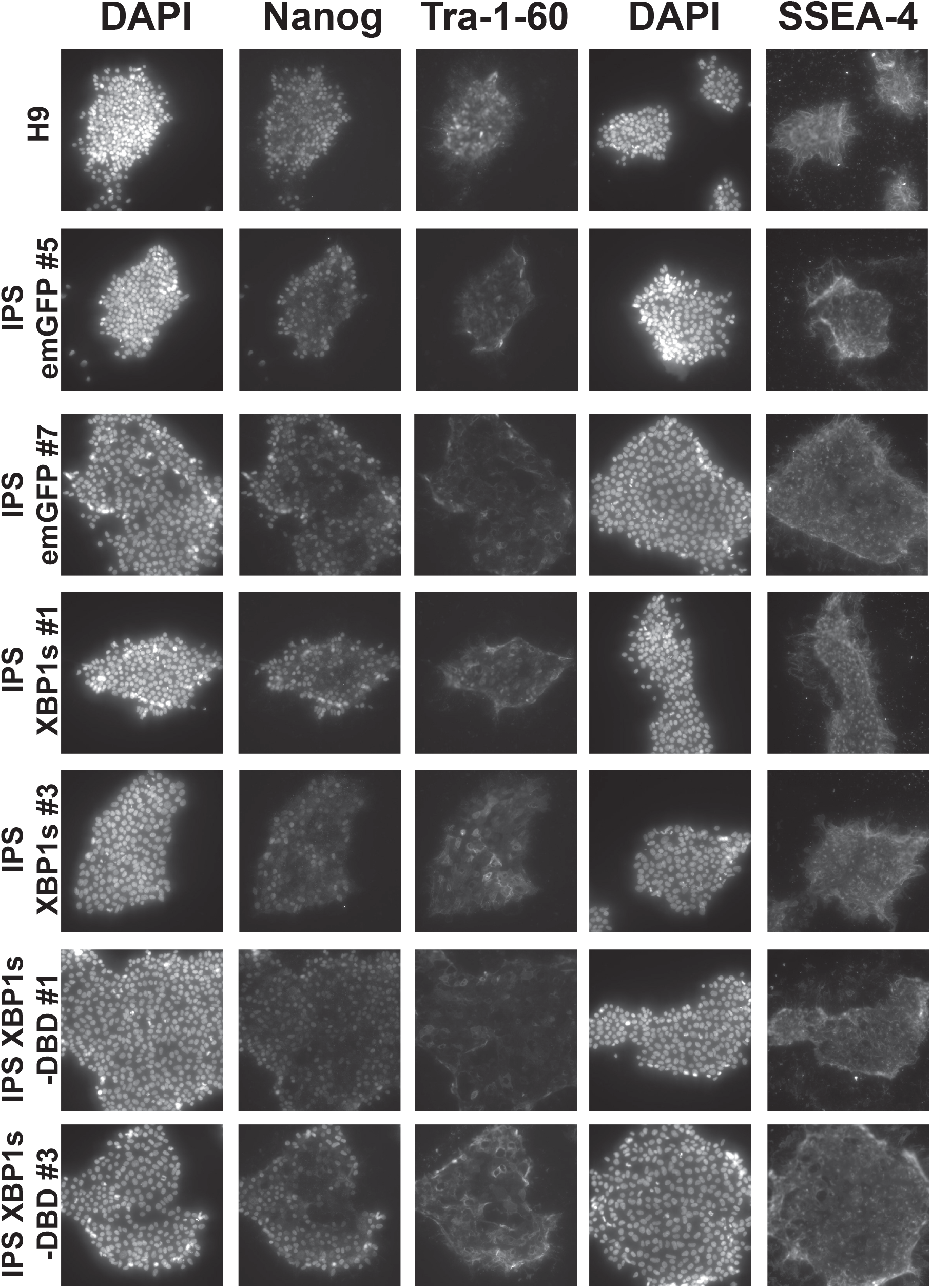
Derived iPSCs stain positive for pluripotent genes. Fluorescent immunostaining of stemness markers Nanog (transcription factor expected localize in the nucleus), TRA-1-60 and SSEA-4 (both cell surface proteins) with DAPI. This was used as binary experiment, the presence of these factors attests of the pluripotency state as the cells they are derived from are negative for them. There was no intent to compare the fluorescence between the cell lines therefore we didn’t provide any quantification. A secondary only control was done and showed no background (data not shown). No scale bar is provided.

**Fig. S6:**
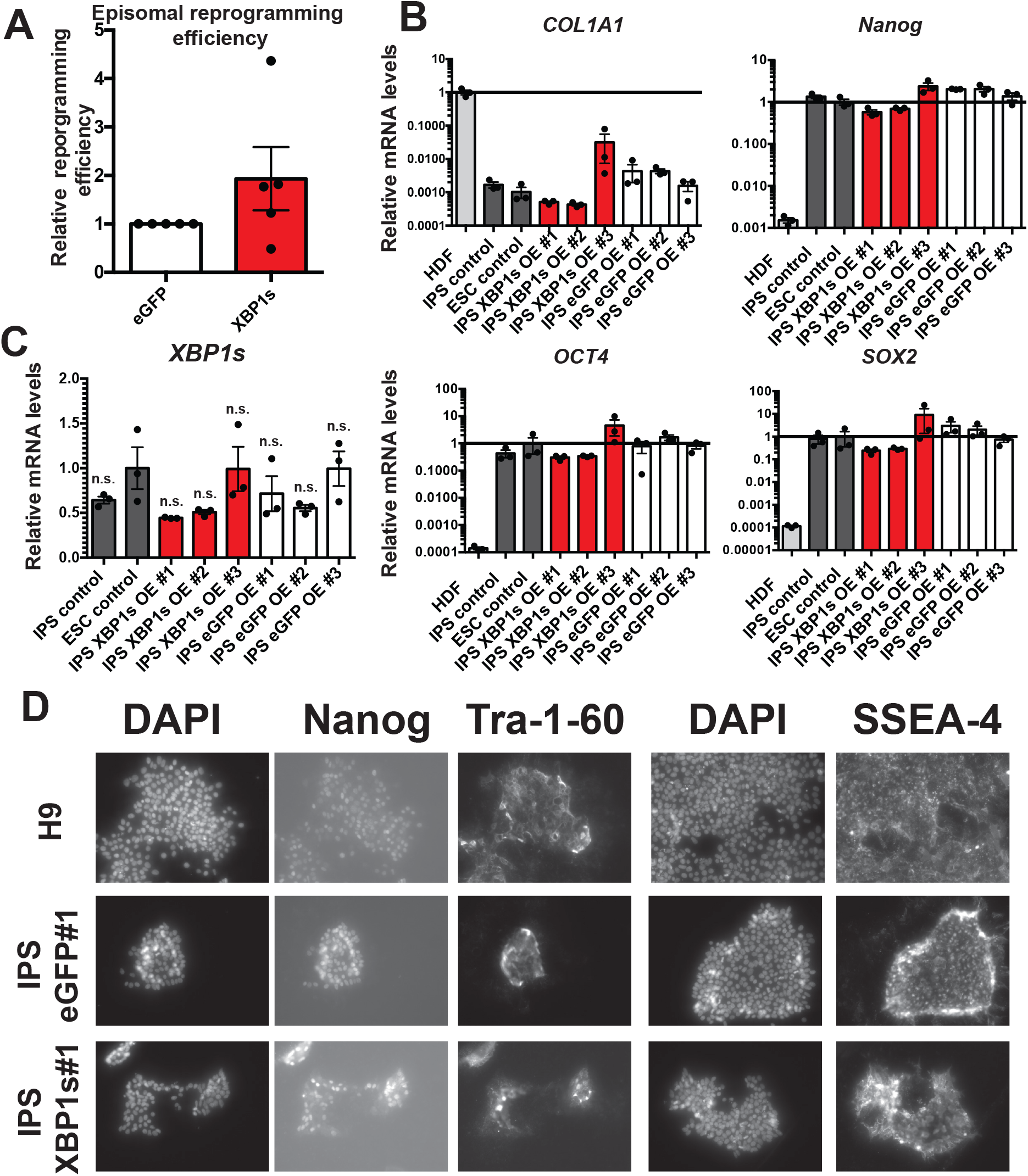
Episomal reprogramming of fibroblasts by XBP1s overexpression. (A) Relative reprogramming efficiency (average +/− SEM). CMV promoter was used to overexpress the transgenes. The differences were not statistically significant using an unpaired two-tailed t-test, n=5. (B) Relative mRNA levels of three stemness markers (*Nanog, SOX2* and *OCT4*) and a fibroblast marker (*COL1A1*) relative to *GAPDH* determined by qRT-PCR (n=3, average +/− SD). H9 line was used as ESC control and iPSC C1 OSKM line (*18*) was used as iPSC control. Values for H9 ESCs were set to 1 for stemness genes while human dermal fibroblast (HDF) values were set to 1 for fibroblast marker *COL1A1.* (C) Relative mRNA levels of *XBP1s* relative to *GAPDH* determined by qRT-PCR (n=3, average +/− SD). n.s. indicates non-significant statistical difference (p-value<0.05) using a Dunnett’s comparison test to control ESC. (D) Fluorescent immunostaining of stemness markers Nanog (transcription factor expected localize in the nucleus), TRA-1-60 and SSEA-4 (both cell surface proteins) with DAPI. This was used as binary experiment, the presence of these factors attests of the pluripotency state as the cells they are derived from are negative for them. There was no intent to compare the fluorescence between the cell lines therefore we didn’t provide any quantification. A secondary only control was done and showed no background (data not shown). No scale bar is provided.

**Fig. S7:**
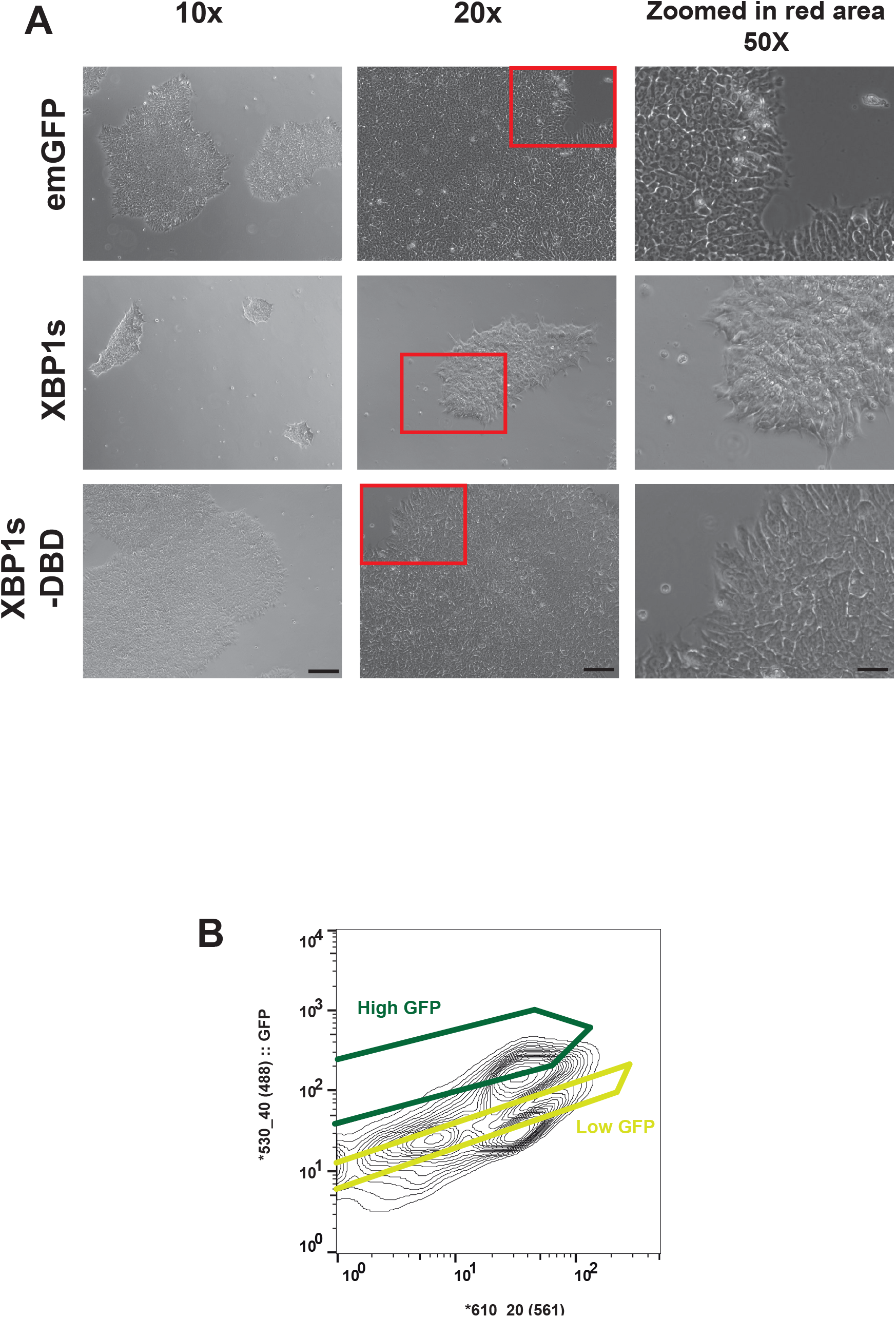
Activation of the UPR^ER^ in stem cells prevents their proper spreading and cell sorting strategy. (A) Morphology of H9 ESC colonies overexpressing *emGFP, XBP1s* or *XBP1s-DBD* driven by EF1α promoter after selection. Scale bar for 10x is 20μm, 10μm for 20X and 4μm for 50X zoomed in. Cells overexpressing *emGFP* or *XBP1s-DBD* have clear nuclei with nucleoli visible while *XBP1s* cells don’t. The latter colonies appear to have a more tridimensional structure. Those cells were kept under puromycin selection for 7 days after tansduction. *XBP1s* cells will eventually be lost. (B) Cells were sorted based on their high/low GFP profile. To take into account cells’ auto-fluorescence, we plotted GFP against 610/20 (561) and gated accordingly.

**Fig. S8:**
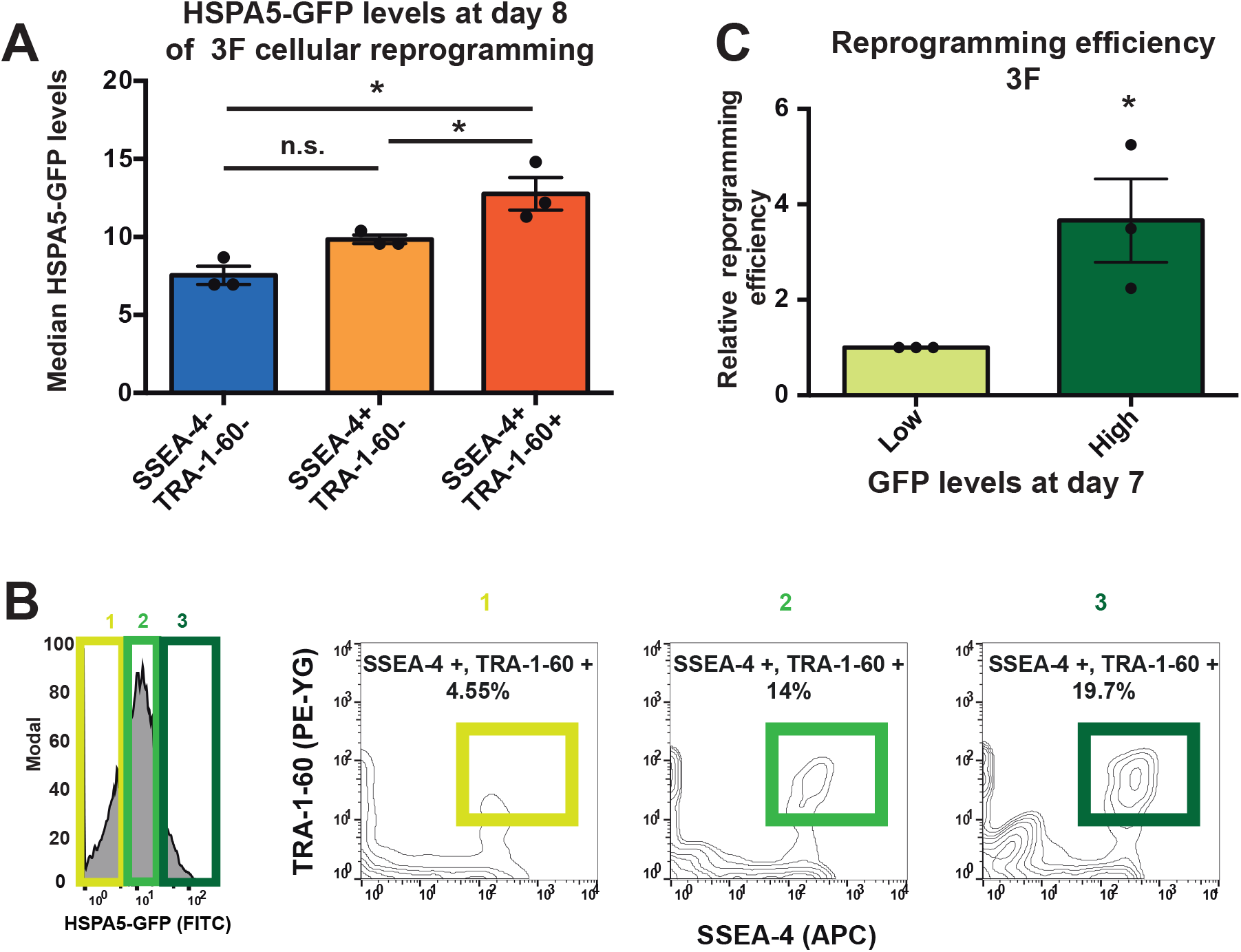
Levels of UPR^ER^ activation are predictive of the reprogramming efficiency using 3F. (A) Median HSPA5-GFP of the different cell states during reprogramming with 3F (n=3, average +/− SD). * indicates statistical significant difference between them (p-value<0.05) using a Newman-Keuls multiple comparison test. (B) Histogram of fibroblast-like HSPA5-GFP at day 8 of reprogramming with 3F. 1, 2, 3 subdivide the population into 3 equal parts. Each of them is represented in the right panel by their SSEA-4 and TRA-1-60 staining. The percentage of double positive cells within each of these populations is shown. (C) Relative reprogramming efficiency of fibroblast-like HSPA5-GFP sorted at day 7 of reprogramming based on their GFP levels and assessed by TRA-1-60 colony staining (n=3, average +/− SEM). * indicates statistical difference (p-value<0.05) using an unpaired two-tailed t-test.

**Table S1:** Transcriptome analysis of UPR^ER^ genes in fibroblasts, iPSCs and ESCs. Transcriptome analysis of UPR^ER^ genes in fibroblasts, IPSCs and ESCs. The data analysis of Lowry and colleague data set (26) was done by Soufi and colleague (35). Here we present a subset of their analysis focusing on UPR effectors. As control, we picked *Nanog* a stemness marker and *COL1A1* a fibroblast marker.

| RefSeq annotation |  |  | Gene expression log2 (GCRMA Intensities) |  |  |
| --- | --- | --- | --- | --- | --- |
| Transcript ID | Gene name | Category | BJ fibroblasts | iPS | ES |
| NM_007348 | *ATF6* | UPR effector | 8.7025125 | 8.090600833 | 7.826710833 |
| NM_182810 | *ATF4* |  | 13.4154 | 13.2858 | 12.74163333 |
| NM_005080 | *XBP1* |  | 8.555195 | 8.339443333 | 8.249261667 |
| NM_005347 | *HSPA5* |  | 11.02015167 | 10.60684667 | 10.482925 |
| NM_003299 | *HSP90B1/GRP94* |  | 11.143225 | 11.47603333 | 11.61726667 |
| NM_014330 | *PPP1R15A/GADD34* |  | 9.644111667 | 8.035643333 | 7.938243333 |
| NM_004083 | *DDIT3/CHOP* |  | 11.4228 | 8.592626667 | 8.161856667 |
| NM_024865 | *NANOG* | Stemness marker | 5.443613333 | 12.3643 | 12.88126667 |
| NM_000088 | *COL1A1* | Fibroblast marker | 12.27041533 | 9.244766 | 10.051136 |

**Table S2:**
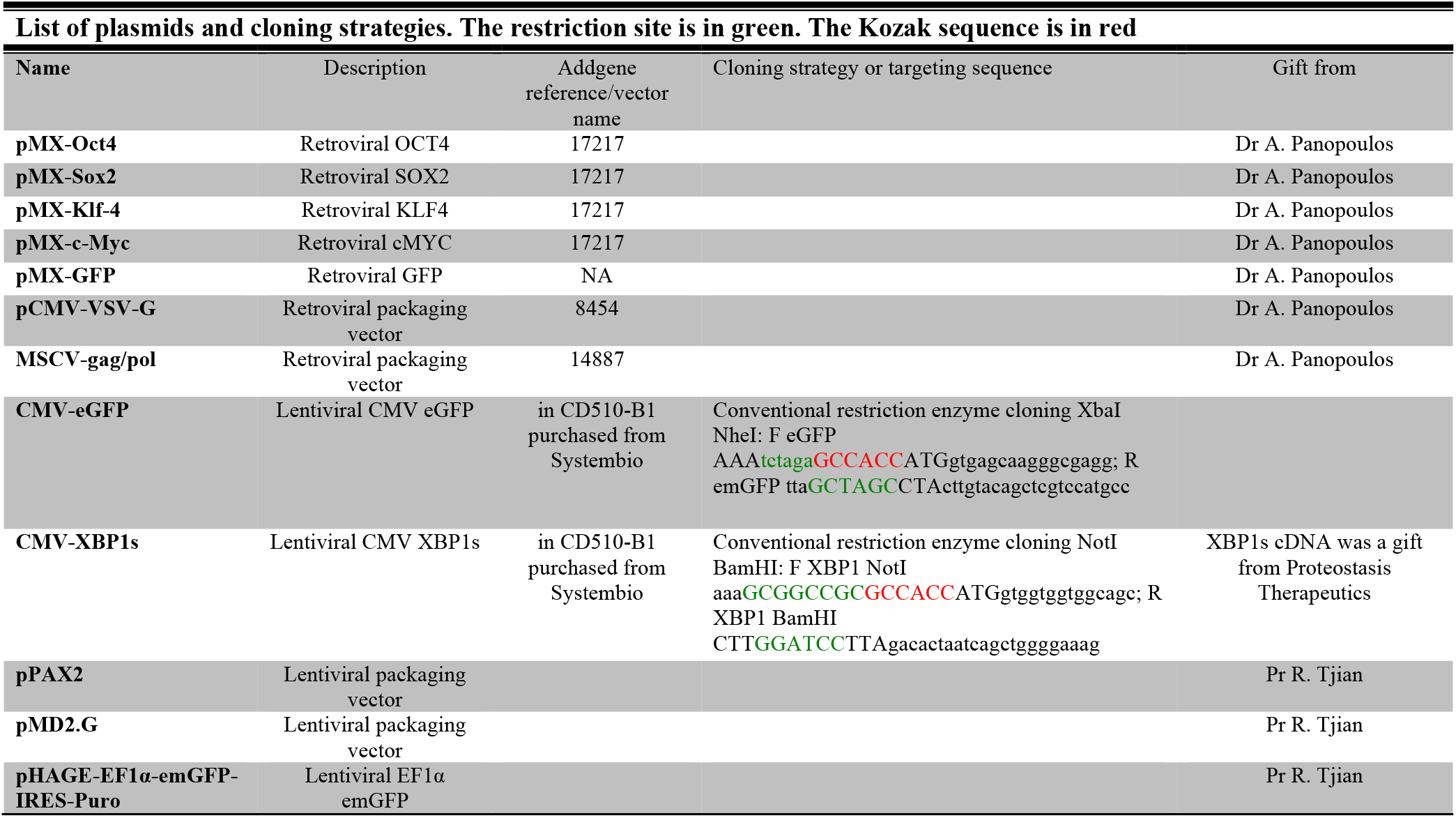
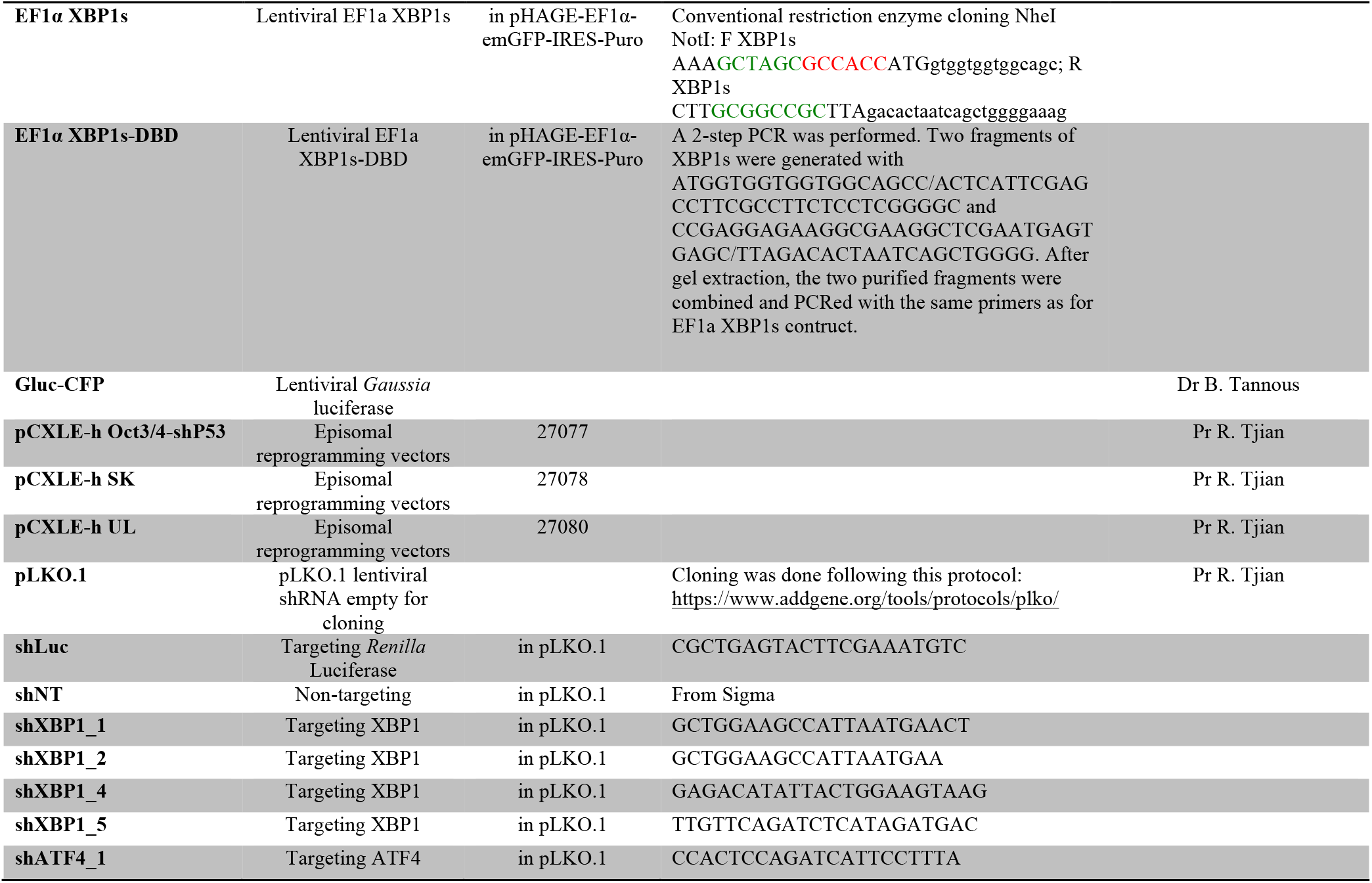
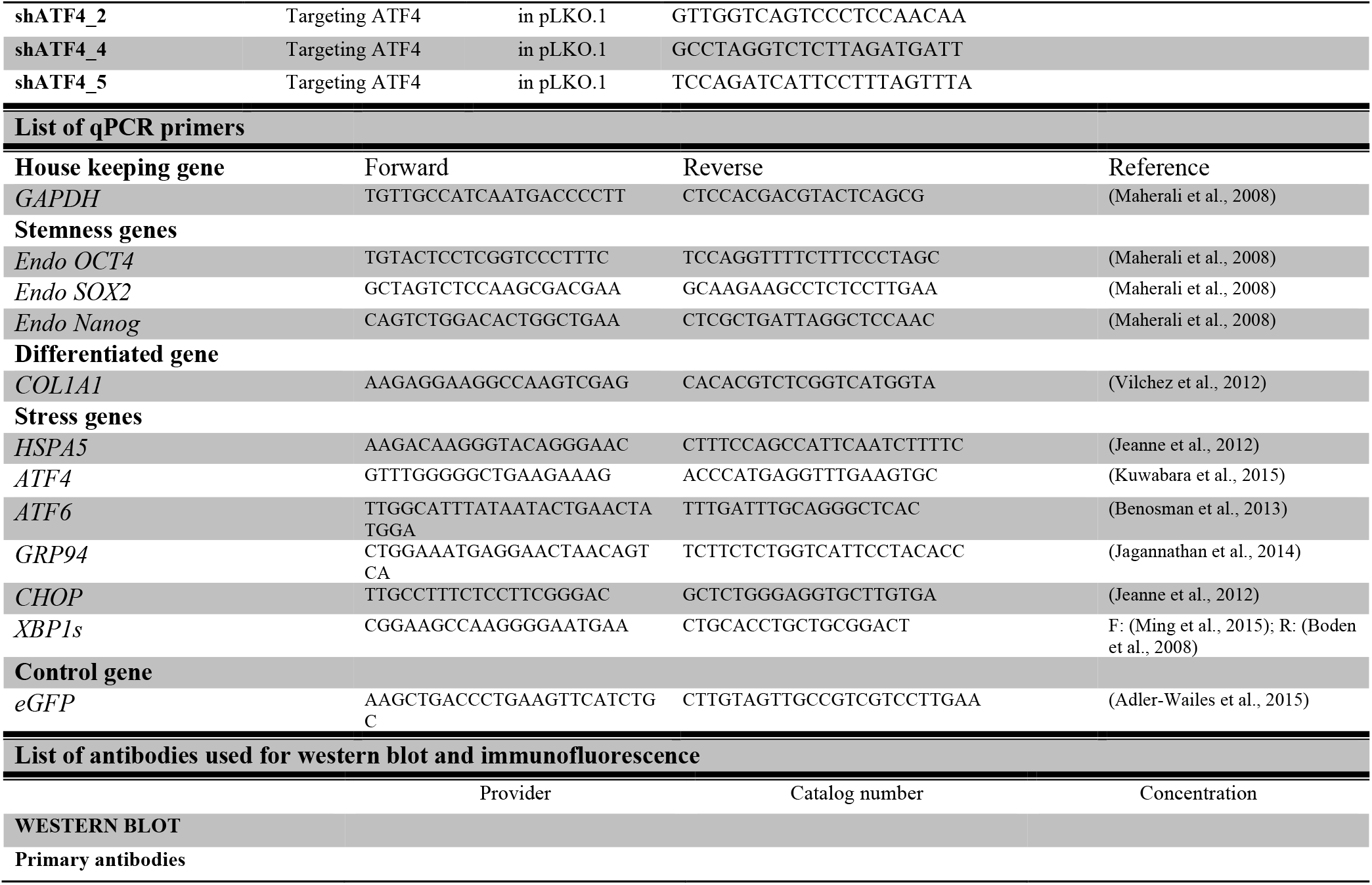
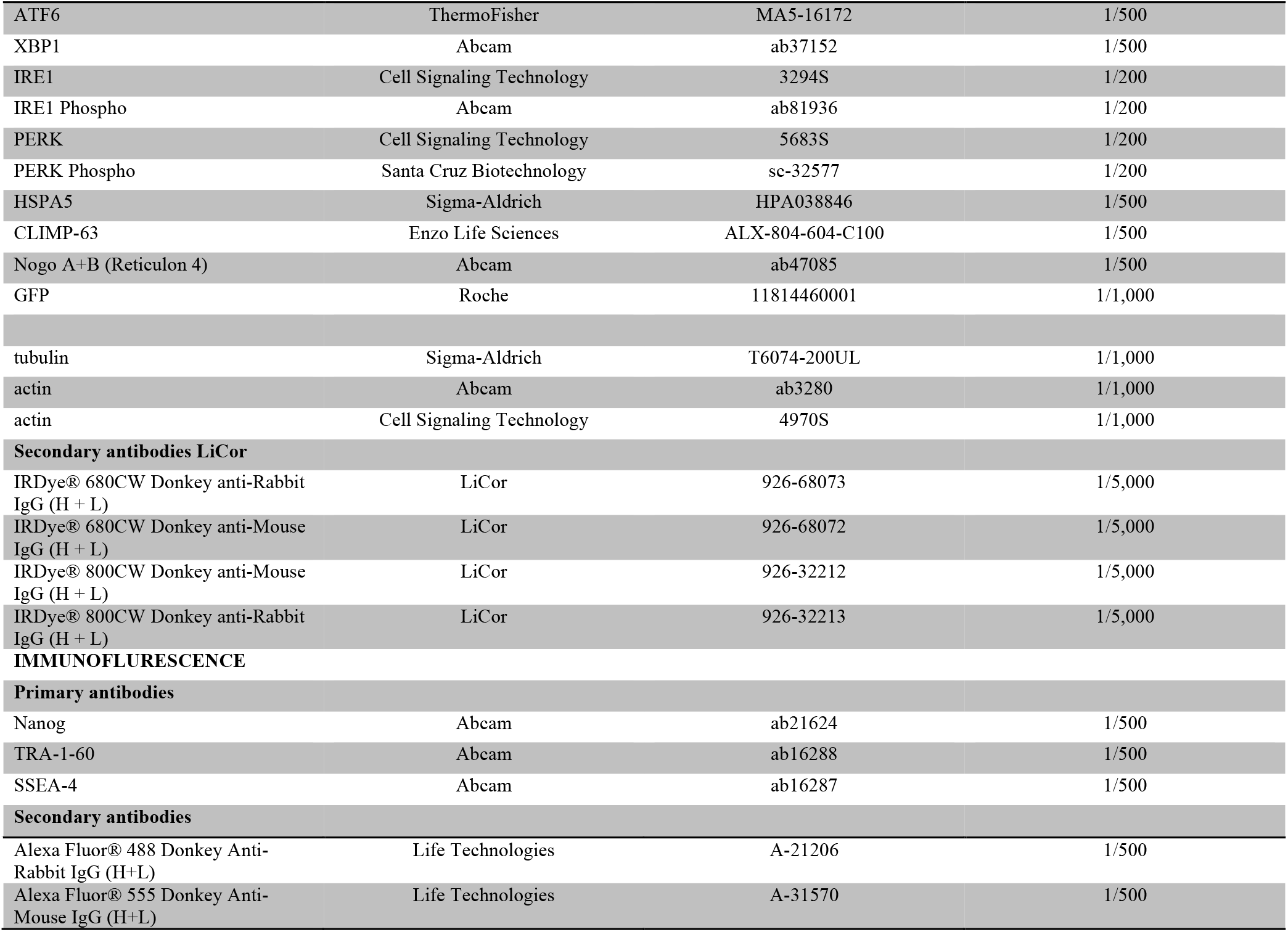
List of reagents used. This table includes the list of plasmids used with the cloning strategy, the list of qPCR primers and the list of antibodies used.

## References

1. J. M. Polo et al., A Molecular Roadmap of Reprogramming Somatic Cells into iPS Cells. Cell. 151, 1617–1632 (2012).

2. K. Takahashi, S. Yamanaka, Induction of Pluripotent Stem Cells from Mouse Embryonic and Adult Fibroblast Cultures by Defined Factors. Cell. 126, 663–676 (2006).

3. Y. Buganim, D. A. Faddah, R. Jaenisch, Mechanisms and models of somatic cell reprogramming. Nat. Rev. Genet. 14, 427–439 (2013).

4. K. Takahashi, S. Yamanaka, A decade of transcription factor-mediated reprogramming to pluripotency. Nat. Rev. Mol. Cell Biol. advance online publication (2016), doi:10.1038/nrm.2016.8.

5. T. Vierbuchen, M. Wernig, Molecular Roadblocks for Cellular Reprogramming. Mol. Cell. 47, 827–838 (2012).

6. D. Vilchez, M. S. Simic, A. Dillin, Proteostasis and aging of stem cells. Trends Cell Biol. 24, 161–170 (2014).

7. P. Walter, D. Ron, The Unfolded Protein Response: From Stress Pathway to Homeostatic Regulation. Science. 334, 1081–1086 (2011).

8. M. Calfon et al., IRE1 couples endoplasmic reticulum load to secretory capacity by processing the XBP-1 mRNA. Nature. 415, 92–96 (2002).

9. H. Yoshida, T. Matsui, A. Yamamoto, T. Okada, K. Mori, XBP1 mRNA Is Induced by ATF6 and Spliced by IRE1 in Response to ER Stress to Produce a Highly Active Transcription Factor. Cell. 107, 881–891 (2001).

10. H. P. Harding, Y. Zhang, D. Ron, Protein translation and folding are coupled by an endoplasmic-reticulum-resident kinase. Nature. 397, 271–274 (1999).

11. K. M. Vattem, R. C. Wek, Reinitiation involving upstream ORFs regulates ATF4 mRNA translation in mammalian cells. Proc. Natl. Acad. Sci. U. S. A. 101, 11269–11274 (2004).

12. K. Haze, H. Yoshida, H. Yanagi, T. Yura, K. Mori, Mammalian transcription factor ATF6 is synthesized as a transmembrane protein and activated by proteolysis in response to endoplasmic reticulum stress. Mol. Biol. Cell. 10, 3787–3799 (1999).

13. Y. Wu et al., Autophagy and mTORC1 regulate the stochastic phase of somatic cell reprogramming. Nat. Cell Biol. 17, 715–725 (2015).

14. K. Kratochvílová et al., The role of the endoplasmic reticulum stress in stemness, pluripotency and development. Eur. J. Cell Biol. 95, 115–123 (2016).

15. T. Namba, T. Ishihara, K. Tanaka, T. Hoshino, T. Mizushima, Transcriptional activation of ATF6 by endoplasmic reticulum stressors. Biochem. Biophys. Res. Commun. 355, 543–548 (2007).

16. J. R. Friedman, G. K. Voeltz, The ER in 3D: a multifunctional dynamic membrane network. Trends Cell Biol. 21, 709–717 (2011).

17. C. E. Badr, J. W. Hewett, X. O. Breakefield, B. A. Tannous, A Highly Sensitive Assay for Monitoring the Secretory Pathway and ER Stress. PLoS ONE. 2, e571 (2007).

18. S. Ruiz et al., Generation of a Drug-inducible Reporter System to Study Cell Reprogramming in Human Cells. J. Biol. Chem. 287, 40767–40778 (2012).

19. T. Brambrink et al., Sequential Expression of Pluripotency Markers during Direct Reprogramming of Mouse Somatic Cells. Cell Stem Cell. 2, 151–159 (2008).

20. E. M. Chan et al., Live cell imaging distinguishes bona fide human iPS cells from partially reprogrammed cells. Nat. Biotechnol. 27, 1033–1037 (2009).

21. D. Hockemeyer et al., A Drug-Inducible System for Direct Reprogramming of Human Somatic Cells to Pluripotency. Cell Stem Cell. 3, 346–353 (2008).

22. T. S. Mikkelsen et al., Dissecting direct reprogramming through integrative genomic analysis. Nature. 454, 49–55 (2008).

23. C. Hetz, E. Chevet, H. P. Harding, Targeting the unfolded protein response in disease. Nat. Rev. Drug Discov. 12, 703–719 (2013).

24. K. Okita et al., A more efficient method to generate integration-free human iPS cells. Nat. Methods. 8, 409–412 (2011).

25. X. Xia, Y. Zhang, C. R. Zieth, S.-C. Zhang, Transgenes Delivered by Lentiviral Vector are Suppressed in Human Embryonic Stem Cells in A Promoter-Dependent Manner. Stem Cells Dev. 16, 167–176 (2007).

26. W. E. Lowry et al., Generation of human induced pluripotent stem cells from dermal fibroblasts. Proc. Natl. Acad. Sci. 105, 2883–2888 (2008).

27. J. Hollien, J. S. Weissman, Decay of Endoplasmic Reticulum-Localized mRNAs During the Unfolded Protein Response. Science. 313, 104–107 (2006).

28. E. Chevet, C. Hetz, A. Samali, Endoplasmic Reticulum Stress–Activated Cell Reprogramming in Oncogenesis. Cancer Discov. (2015), doi:10.1158/2159-8290.CD-14-1490.

29. D. Acosta-Alvear et al., XBP1 Controls Diverse Cell Type- and Condition-Specific Transcriptional Regulatory Networks. Mol. Cell. 27, 53–66 (2007).

30. D. Vilchez et al., Increased proteasome activity in human embryonic stem cells is regulated by PSMD11. Nature. 489, 304–308 (2012).

31. S. M. Buckley et al., Regulation of Pluripotency and Cellular Reprogramming by the Ubiquitin-Proteasome System. Cell Stem Cell. 11, 783–798 (2012).

32. T. T. Onder et al., Chromatin-modifying enzymes as modulators of reprogramming. Nature. 483, 598–602 (2012).

33. D. Hockemeyer et al., Genetic engineering of human pluripotent cells using TALE nucleases. Nat. Biotechnol. 29, 731–734 (2011).

34. H.-E. Kim et al., Lipid Biosynthesis Coordinates a Mitochondrial-to-Cytosolic Stress Response. Cell. 166, 1539–1552.e16 (2016).

35. A. Soufi, G. Donahue, K. S. Zaret, Facilitators and Impediments of the Pluripotency Reprogramming Factors’ Initial Engagement with the Genome. Cell. 151, 994–1004 (2012).

